# 14-3-3 shuttles Activity-dependent neuroprotective protein to the cytoplasm to promote appropriate neuronal morphogenesis, cortical connectivity and calcium signaling

**DOI:** 10.1101/2020.05.26.105015

**Authors:** Sarah A. Bennison, Sara M. Blazejewski, Xiaonan Liu, Kazuhito Toyo-oka

**Affiliations:** Department of Neurobiology and Anatomy, Drexel University College of Medicine Department of Pharmacology and Physiology, Drexel University College of Medicine Philadelphia, PA 19129 USA

## Abstract

Neurite formation is the earliest stage of neuronal morphogenesis, where primitive dendrites and the primitive axon emerge from a spherical neuron and begin to elongate. Defective neuritogenesis is a contributing pathogenic mechanism behind a variety of neurodevelopmental disorders. Activity-dependent neuroprotective protein (Adnp) is essential to embryonic and postnatal brain development, and mutations in *ADNP* are among the most frequent underlying autism spectrum disorder (ASD). We found that knockdown of Adnp *in vitro* and *in vivo* in mouse layer 2/3 pyramidal neurons leads to increased neurite initiation and defective neurite elongation, suggesting that Adnp has distinct roles in each. *In vivo* analysis revealed that deficits begin at P0 and are sustained throughout development, the most notable of which include increased neurite stabilization, disrupted angle of the apical dendrite, increased basal dendrite number, and increased axon length. Because small changes in neuronal morphology can have large-scale effects on neuronal function and connectivity, we performed *ex vivo* calcium imaging to assess spontaneous function of layer 2/3 pyramidal neurons deficient in Adnp. This revealed that Adnp deficient neurons had a greater spontaneous calcium influx and a higher proportion of cells firing action potentials. Next, we utilized GRAPHIC, a novel synaptic tracing technology, to assess interhemispheric cortical connectivity. We found increased interhemispheric excitatory connectivity between Adnp deficient layer 2/3 pyramidal neurons. Because Adnp is a multifunctional protein with both transcription factor and cytoskeletal activity, we performed localization analysis of Adnp as neurons underwent neurite formation to probe the mechanism of our morphological defects. We found that Adnp is shuttled from the nucleus to the cytoplasm upon differentiation and this shuttling can be blocked via application of a global 14-3-3 inhibitor, difopein. Furthermore, we found that Adnp binds nuclear-cytoplasmic shuttle 14-3-3ε. We conclude that Adnp is shuttled from the nucleus to the cytoplasm by 14-3-3ε, where it regulates neuronal morphology, maturation, cortical connectivity, and calcium signaling.

## Introduction

Neuritogenesis is a fundamental step of cortical development essential for establishing correct neuronal morphology, connectivity, and function (1, 2). Immature neurons have a spherical morphology that upon maturation develops to extend a single axon and multiple dendrites (3–6). This complex feat is driven by neuronal polarization which causes an initial break in symmetry within an immature neuron, seamlessly followed by the two stages of neuritogenesis: neurite initiation followed by neurite elongation (4, 7, 8). Following neuronal polarization, actin aggregates form the sites of primitive neurites, precursors to the axon and dendrites (4, 9, 10). At these sites, actin rich filopodia and lamellipodia rapidly extend and retract before being stabilized by microtubule invasion. Microtubules then drive neurite elongation as primitive neurites are lead to their appropriate destinations where they differentiate into a single axon and multiple dendrites (8, 11–14). It is of current interest which proteins are key players in this early developmental process, particularly those which are causatively mutated in neurodevelopmental disorders.

Following neurite formation, dendrites mature and develop intricately branched, complex structures that form synaptic contacts with neighboring neurons that, when formed correctly, lead to functional connectivity (1, 2, 15). The sites of the vast majority of excitatory connections are through dendritic spines, the number, developmental timing, and morphology of which are crucial to form appropriate functional connections (16, 17). These early steps in cortical development are essentially integrated and complex, with each stage relying on the fidelity of the others. When one or more of these stages go awry the result is a variety of neurodevelopmental disorders such as autism spectrum disorder (ASD), intellectual disability (ID), schizophrenia, and Down syndrome (8, 18–21). Thorough investigations of dendritic maturation and synaptic connectivity in connection with the etiology of many neurodevelopmental diseases have been performed, however earlier stages of neuronal morphology such as neuritogenesis have yet to be elucidated. Defective neuritogenesis has been observed and strongly implicated as a contributing pathogenic factor in many models of neurodevelopmental diseases, but a mechanistic understanding as to how and why neurite formation goes awry in disease states is yet to be understood.

Activity-dependent neuroprotective protein (Adnp) is highly conserved with extremely diverse functions essential in both the central nervous system and throughout the body (22). Mutations in *ADNP* are well characterized as some of the most frequent underlying ASD and intellectual disability (ID) and lead to a neurodevelopmental disorder known as ADNP syndrome (22, 23). Symptoms of the ADNP syndrome range from moderate to severe and the hallmarks include ID, ASD, delayed speech and motor development, sleep disorder, and seizures (22–28). Patients also have disorders of multiple organ systems including the digestive system and heart abnormalities (23) and may be characterized/diagnosed by early tooth eruption (26). However, there are currently no approved treatment options for patients with mutations in ADNP, rendering a greater understanding of ADNP’s functions during development and mutational etiology of high importance. Furthermore, deficits in ADNP have also been associated with neurodegeneration and Tau pathology. A recent paper identified somatic mutations in *ADNP* driving Tau-microtubule dissociation and increased tauopathy (25), in line with early discoveries of tauopathy in an animal model of ADNP deficiency (29).

Adnp interacts with microtubules to promote polymerization through an 8-amino acid sequence referred to as “NAP” (22, 29). Treatment of cells in culture with NAP has been shown to promote or rescue neurite outgrowth (30, 31), enhance microtubule dynamics and Tau-microtubule association (29, 32, 33), and interact with EB3 to enhance dendritic spine formation (34, 35). Taken together, these results position Adnp as an ideal candidate to regulate neurite formation during development. However, a detailed explanation of Adnp’s role in this neuronal process has yet to be elucidated and an *in vivo* neuritogenesis analysis has yet to be performed. Failure of neurite outgrowth or degeneration of neurites underlies a variety of neurodevelopmental disorders many of which share symptomology with ADNP syndrome (8, 20, 21, 23). Rescue of neurite formation defects, specifically by NAP, has also proved a potentially valuable therapeutic avenue for a variety of disorders (30, 36). However, the details of Adnp’s involvement in neurite formation *in vivo* and whether defective neurite formation plays a pathogenic role in ADNP syndrome during this early foundational stage of cortical development has yet to be elucidated.

The purpose of this study was to perform a detailed, multi-level analysis of Adnp’s functions in neurite formation. We performed in-depth *in vitro* and *in vivo* analyses of the consequences of knockdown of Adnp in layer 2/3 pyramidal neurons in the somatosensory cortex on multiple stages of cortical development, with a focus on neuritogenesis. Surprisingly, Adnp’s role in neurite formation is not as clear as previously suggested. We show that knockdown of Adnp in layer 2/3 pyramidal neurons leads to an increase in neurite number, increase in the length of the axon, but decrease in length of the basal dendrites. A further developmental defect discovered was disruption of the angle of the apical dendrite, potentially effecting functional connectivity. *Ex vivo* time-lapse live imaging of neuritogenesis at P0 corroborated our fixed analyses, as well as revealed further defects involving dilations and swellings of growing neurites and defective growth speed. We also noted a defect previously reported in the Adnp haploinsufficient mouse of decreased dendritic spine density (34). We further quantified dendritic spine morphology and found that Adnp knockdown neurons had more immature spines. Functionally, we uncovered increased spontaneous calcium signaling and interhemispheric cortical connectivity in Adnp deficient pyramidal neurons through excitatory shaft synapses, potentially negating reported dendritic spine defects.

As previously implied (37, 38), we found that Adnp expression changes in subcellular localization as primary cortical neurons undergo differentiation from neuronal precursor cells to mature neurons, beginning exclusively in the nucleus and traveling to the cytoplasm as development progresses. This suggests that Adnp’s role in neuronal morphogenesis may be mainly due to Adnp’s cytoplasmic activities, instead of its other known functions as a transcription factor. Most importantly, we identified 14-3-3ε as a candidate molecular shuttle. These studies are the first to perform such a detailed analysis of the consequences of loss of Adnp on neurite formation and neuronal morphology, revealing a wealth of knowledge on the ways reduced expression of Adnp effects such crucial stages of cortical development, and providing new insights into the ways in which mutations in Adnp may result in pathology.

## Methods

### Mice

C57BL/6 mice were maintained in house, and females and males were used for *in utero* electroporation (IUE) and primary neuronal culture unless otherwise described. All experimental procedures were approved by the Institutional Animal Care and Use Committee of Drexel University. The day of the detection of the vaginal plug was defined as embryonic (E) 0.5.

### Plasmids

Mouse Adnp cDNA (pENTR223.1-Adnp, BC167195) and mammalian expression vector (pCMV6-Entry-Adnp-Myc-DDK, MR223066) were purchased from TransOMIC and Origene, respectively. Mouse Adnp was amplified from pENTR223.1-Adnp using Q5 High Fidelity DNA Polymerase (NEB) and primers containing 6xHis tag to insert it into the C-terminal region of Adnp, and the PCR fragment was cloned into pLV-CAG1.1-P2A-mScarlet plasmid, which was created by inserting P2A-mScarlet fragment into pLV-CAG1.1 plasmid (kind gift from Dr. Masahito Ikawa in Osaka University, Japan), to create pLV-CAG1.1-Adnp-6xHis-P2A-mScarlet. Adnp shRNA was designed using the web-based design tools, siRNA Wizard Software (InvivoGen) and BLOCK-iT RNAi Designer (ThermoFisher Scientific), and oligos were synthesized by IDT. The annealed oligos were cloned into pSCV2-Venus, mScarlet, and mTagBFP2 plasmids (Hand and Polleux, 2011, Wachi, et al., 2015). pSCV2-mScarlet and pSCV2-mTagBFP2 were created by replacing Venus in pSCV2-Venus into mScarlet and mTagBFP2 by PCR. The target sequence was GAGCCTGTACCGAAGGTTA. Scramble shRNA (ACTACCGTTGTTATAGGTG) was used as a negative control. shRNA-resistant Adnp was created by PCR using primers in which 6 nucleotides were mutated. The shRNA-resistant Adnp sequence is GA<u>A</u>CC<u>A</u>GT<u>T</u>CC<u>C</u>AA<u>A</u>GT<u>A</u>A. pEYFP-difopein, 14-3-3 peptide inhibitor, is a kind gift from Dr. Yi Zhou at Florida State University.

### Antibodies

Primary antibodies used in this studies were as follows: Anti-ADNP (Rabbit, Abm, Y409055), Anti-ADNP antibody (Mouse, F-5, Santa Cruz Technology, sc-393377), Anti-Sox2 (Goat, Y-17, Santa Cruz Technology, sc-17320), Anti-type III β-tubulin (Mouse, 2G10, ThermoFisher Scientific, MA1-118), Anti-MAP2 (Mouse, HM-2, Sigma, M4403), Anti-GAPDH (Mouse, Proteintech, 60004-1-Ig), Anti-His-tag antibody (mouse, Proteintech, 66005-1-Ig), Anti-HA-tag antibody (mouse, 12CA5, Roche, 11583816001), Anti-Brn2 antibody (Rabbit, Proteintech, 14596-1-AP), The following secondary antibodies were used: FITC-conjugated Donkey-anti-Rabbit IgG (Jackson ImmunoResearch Laboratories, 711-096-152), FITC-conjugated Donkey-anti-Goat IgG (Jackson ImmunoResearch Laboratories, 705-095-147), TRITC-conjugated Donkey-anti-Mouse (Jackson ImmunoResearch Laboratories, 715-025-151), Cy5-conjugated Donkey-anti-Mouse (Jackson ImmunoResearch Laboratories, 715-175-150), and Cy5-conjugated Donkey-anti-Rabbit (Jackson ImmunoResearch Laboratories, 711-175-152). Fluorescent western blot was performed using IRDye 680 RD Donkey-anti-mouse (LI-COR, 926-68072).

### Primary Neuron and Neurosphere Culture

Primary cortical neurons were harvested from E15.5 mouse embryonic cortices for mature neuron culture and E14.5 for neurosphere culture. Briefly, pregnant dams were euthanized using CO2 and the embryos were immediately removed and decapitated in ice-cold phosphate-buffered saline. Using a dissection microscope, cortices were harvested from embryonic brains and placed in phosphate-buffered saline on ice. Cortices were dissociated using 0.01% trypsin and a 1000*μ*L pipette to mechanically dissociate cortices into a single cell suspension. To inactivate trypsin, 200*μ*L of 50*μ*g/mL bovine serum albumin (BSA) was added. Cells were passed through a filter to remove excess debris, washed with phosphate-buffered saline and centrifuged for 5 minutes. This process was repeated twice, and cells were counted. To introduce genetic constructs into neurons, we placed 3-5 million cells into cuvettes for nucleofection (Amaxa). 10*μ*g of DNA was added per cuvette. Cells were then plated in Neurobasal media supplemented with B27 for mature neurons or DMEM/F12 supplemented with 1% bFGF and EGF for neurospheres. For neurospheres, media was changed every 3 days and cells were kept in culture for 14 days. Mature neurons were re-plated onto coverglass 48 hours following nucleofection, once the transfection had reached its peak effect, and were allowed to grow for 48 more hours until neurite elongation had proceeded to completion.

### Histology and Immunofluorescence Staining

To analyze Adnp expression, brains were dissected at postnatal day (P)15 and fixed with 4% paraformaldehyde/Phosphate-buffered saline overnight at 4°C. Fixed samples were cryo-protected by addition of 25% sucrose/Phosphate-buffered saline for 48 hours at 4°C. Samples were embedded with O.C.T. compound (Sakura) and stored at −80°C. Cryo-sectioning (60 µm thickness) was performed by cryostat (Micron HM505 N) and slices were air-dried. Sections were rinsed three times in Tris-buffered saline and treated with 0.2% Triton X-100/Tris-buffered saline for 10 minutes at room temperature, followed by blocking for 30 minutes in 5% Bovine serum albumin/Phosphate-buffered saline supplemented with 0.25% Tween-20 to prevent nonspecific binding. Primary antibodies were diluted in blocking buffer, and sections were incubated in primary antibody overnight at 4°C. Secondary antibodies were diluted with blocking buffer and sections were incubated for 30 minutes at room temperature. Sections were stained with 40,6-Diamidino-2-phenylindole, Dihydrochloride (DAPI, 600nM) and embedded with 90% glycerol made with Tris-buffered saline. To validate layer targeting of IUE, brains were dissected at P15 and underwent the same staining protocol.

Neurospheres were stained as described with a few modifications (39). Briefly, neurospheres were cultured for 14 days and transferred to a 15 mL tube by a 1000*μ*L pipette. Spheres were allowed to settle to the bottom of the tube by gravity for 5 minutes before media was removed and spheres were washed with phosphate-buffered saline. This was repeated 3 times before treating with 4% paraformaldehyde/Phosphate-buffered saline for 20 minutes. Neurospheres were then stained using the same protocol for brains but remained free-floating in the 15 mL tube. Finally, neurospheres were stained with 40,6-Diamidino-2-phenylindole, Dihydrochloride (DAPI, 600nM) and embedded with 90% glycerol made with Tris-buffered saline.

Primary neurons for imaging were grown on glass coverslips for imaging. At the time of fixation, neurons were washed three times with phosphate-buffered saline and treated with 4% paraformaldehyde/Phosphate-buffered saline for ten minutes. Cells were immunofluorescently stained using the same protocol for brain slice and neurosphere staining.

All imaging was performed using a confocal microscope (Leica SP8).

### In Utero Electroporation (IUE)

E15.5 pregnant mice were used for IUE as previously described (8, 40, 41). Briefly, under anesthesia, the uterine horn was exposed and 1-2μl of plasmids (1μg/μl) was injected into the lateral ventricle by pulled-glass micropipette. Then, electric pulses (three pulses of 32V) were given by the tweezers-type electrodes over the uterine muscle using CUY21 electroporator (Nepa GENE). The uterine horn was returned into the abdomen, and brain samples were collected at P0, P3 and P15 for analysis.

### Analysis of Neuronal Morphology

*In vitro* and *in vivo,* neurites were classified as cell protrusions from the cell soma greater than 5*μ*m long. To analyze neuronal morphology *in vivo*, brains were dissected at P3 or P15 and fixed with paraformaldehyde/Phosphate-buffered saline overnight at 4°C. Fixed samples were processed as described above. Cryo-sections (60 µm thickness) were cut and stained by DAPI. Imaging was performed using a confocal microscope (SP8 Leica). All image analysis was performed using Fiji software. To analyze the angle at which the apical dendrite extended with respect to the cortical plate, a 90° angle was drawn from the center of the cell soma to the cortical plate using the angle tool. Then, an angle was drawn to the center of the apical dendrite and measured. Polar histograms were generated using MATLAB and depict the angles at which the apical dendrites extended with respect to the cortical plate. The average deviation from the expected 90° was also measured and compared. Number of basal dendrites were counted, and the length of basal and apical dendrites were measured using Fiji. Sholl analysis was performed using the Sholl Analysis Plugin (Gosh Lab, UCSD) for Fiji following the developer instructions. To measure axon length *in vivo*, the length of the axon bundle across the midline was measured using ten brain slices per group of the same brain area. Dendritic spines were characterized based on geometric characteristics previously defined (42). Spines longer than 2µm were classified as filopodia, spines between 1 and 2µm long were classified as long thin, and spines shorter than 1µm were classified as thin. Stubby spines were classified if the length to width ratio was less than 1. Mushroom spines were classified if the spine head width was greater than 0.6µm. Spines with two heads were classified as branched. Standard deviation projection images from z-projection photos produced from z-stack data were used for analysis.

### Fluorescence Analysis

All fluorescence quantification was performed using Fiji and a modified protocol from the Queensland Brain Institute imaging facility (43). Briefly, all fluorescence values are corrected for background contribution and for the area of the cell, to prevent confounds such as larger cells intrinsically having a greater fluorescence intensity value. For each cell, 3 background measurements were taken surrounding the cell. Those measurements were averaged for use in future calculations. A region of interest (ROI) was traced around either the whole cell or different cellular compartments depending on the analysis, and the measurement tool in Fiji was used to extract following measurements: integrated fluorescence density of ROI, area of ROI, and background fluorescence. The following calculation was used to perform the appropriate corrections and obtain the final “corrected fluorescence” value used for analysis and reported in figures: corrected fluorescence = integrated fluorescence density of ROI – (area of ROI x mean fluorescence of background readings).

### Ex Vivo Live Imaging

Brain slice preparation and time-lapse live imaging on brain slices were performed as previously described (8, 40). Briefly, P0 brains were removed and placed in ice-cold artificial cerebrospinal fluid (CSF). Brains were embedded in 4% low-melting agarose and slices were cut with a 300*μ*m thickness in ice cold artificial CSF using a VTS1000 vibratome (Leica). Slices were incubated in D-MEM/F-12 imaging media without phenol red supplemented with 10% FBS for at least 1h at 37 °C, 5% CO2 for recovery. Slices were transferred to a 35mm dish and submerged in neutralized rat tail collagen I (Life Technologies). Collagen solidified for 30 min at 37 °C, 5% CO2. Slices were then covered with imaging media. Time-lapse live imaging was performed using an upright confocal laser scanning microscope (TCS SP2 VIS/405, Leica) with a 20X HCX APO L water-dipping objective (NA 0.5). During imaging, slices were cultured in the imaging media and kept at 37 °C with 95% air/5% CO2 in a stage top chamber incubator (DH-40iL, Warner Instruments). Confocal Z-stack images were taken every 10min for more than 10 hours.

### Calcium Imaging

Mice underwent IUE at E15.5 and brain slice preparation was performed as described for *ex vivo* live imaging with a few modifications using brains of 2-month-old female mice. Slices were incubated in D-MEM/F-12 imaging media without phenol red supplemented with 10% FBS for at least 20 minutes at 37 °C, 5% CO2 for recovery. Slices were transferred into glass-bottom 35mm dishes (MatTek) for imaging. A membrane was placed on top of the slices to reduce movement during imaging. Time-lapse live imaging was performed using an inverted fluorescent microscope (Ziess, Axio Observer Z1) with a 20x objective. Images were captured using Camera Streaming mode set to 1500 cycles, allowing a frame to be taken approximately every 30 ms for 1 minute with 4×4 binning while the slices were maintained at 37°C with a stage top incubator (Zeiss).

All cells included in the analysis were double-positive for GCaMP6s and either Adnp-shRNA-mScarlet or Scramble-shRNA-mScarlet. Image analysis was performed using Zen 2 Pro analysis software (Ziess 2011). Circular regions of interest were placed on the cell soma. Baseline fluorescence (F0) was obtained by averaging the fluorescence intensity inside the region of interest throughout the time course imaging. Fluorescence intensity for the time course was measured by averaging all pixels in the region of interest at each frame of the imaging (Fmeasured). Percent change in fluorescence (ΔF/F0) was calculated as (Fmeasured-F0)/ F0 x 100 for every frame of the time course (44).

### GRAPHIC

IUE was performed to inject two complementary GRAPHIC plasmids (pCAGGS-nGRAPHIC-2A-H2B-mCherry and pCAGGS-cGRAPHIC-T2A-mCherry) and Adnp-shRNA-mTagBFP2 or Scramble-shRNA-mTagBFP2 into the right and left lateral ventricles, respectively, of E15.5 embryos. Transfected cells in the right hemisphere are H2B+, BFP2+ and have GFP puncta where contacts are formed from the opposing hemisphere; the left hemisphere has cells transfected with mCherry, BFP2, and have GFP puncta where contacts are formed from the opposing hemisphere. Embryos developed until P30 when female brains were harvested, cryopreserved, sectioned and imaged as described in the histology methods section. Only mCherry+ and BFP2+ cells were used for analysis, and only GFP puncta co-localized with dendrites were counted.

### Pull-down

COS-1 cells were transfected with either 6xHis-Adnp + HAHA or 6xHis-Adnp + HAHA-14-3-3ε and grown to confluency on 10cm dishes. Cells were collected and pull-down was performed using ant-HA antibody-coated beads (HA-probe (F-70), Santa Crus Biotechnology, sc-7392). Following pull-down, western blot was performed using anti-His antibody to detect 6xHis-Adnp. Also, for input, whole protein lysates before performing the pull-down were used to detect 6xHis-Adnp and HAHA-14-3-3ε.

### Statistical Analysis

The experimenter was blinded during all data acquisition and analysis. All experiments have biological and technical replicates that include performing three experiments and plating duplicates per condition for culture experiments. For IUE, at least 3 brains from two litters were analyzed. Quantitative data were subjected to statistical analysis using SPSS (IBM Analytics) and MATLAB (MathWorks). Data were tested for normality using Shapiro-Wilk’s test with a cut-off of p<0.05. Outliers were removed if their z-score fell outside +/− 2.5 from the mean. The data were analyzed by two-tailed independent-samples t-tests, one-way or two-way ANOVAs with post-hoc test if needed, and chi-squared tests where appropriate. Results from parametric tests were considered significant if p<0.05. All data are presented as mean ± standard error of the mean. Significance is reported on figures as follows: *p<0.05, **p<0.01, ***p<0.001.

## Results

### Adnp knockdown disrupts cortical neuritogenesis in vitro

Previous studies have shown that knockdown of Adnp leads to a decrease in MAP2 fluorescence intensity in a differentiated embryonic carcinoma cell line, P19 cells, suggesting decreased neurite formation (37). Another study has shown that treating primary cortical neurons with NAP increases neurite formation (31). Taken together, these studies implicate Adnp as an important regulator of neurite formation. However, Adnp has differing roles based on both cell-type and developmental timing. Therefore, it is crucial to perform a direct analysis of how Adnp knockdown effects neuritogenesis in cortical neurons. Before moving to our neuronal cell model, we confirmed that Adnp is expressed in primary cortical neurons using immunofluorescence staining (Fig. 1A). We found that Adnp is expressed in both the cell body and along developing neurites (Fig. 1B), with increased concentration in the perinuclear region and at the base of growing neurites. We also confirmed that Adnp is expressed throughout the mouse cortex at P15 and its expression is predominantly in the cytoplasm (Fig. 1C-D), as implied by previous work (37, 38, 45). We harvested primary cortical neurons from E15.5 mouse embryos and introduced Adnp short hairpin RNA (shRNA) or scramble shRNA with a Venus fluorophore via nucleofection. We designed Adnp shRNA not to target any other known sequence in the mouse genome and found it to be >97% efficient at knockdown of Adnp validated by Western blot and by immunofluorescence staining (Supplemental Fig. 1A-D). Scramble shRNA was used as a control and does not target any known sequence in the mouse genome, shares a backbone vector with Adnp shRNA, and was validated by Western blot and immunofluorescence staining of Adnp (Supplemental Fig. 1 A-D).

**Figure 1.**
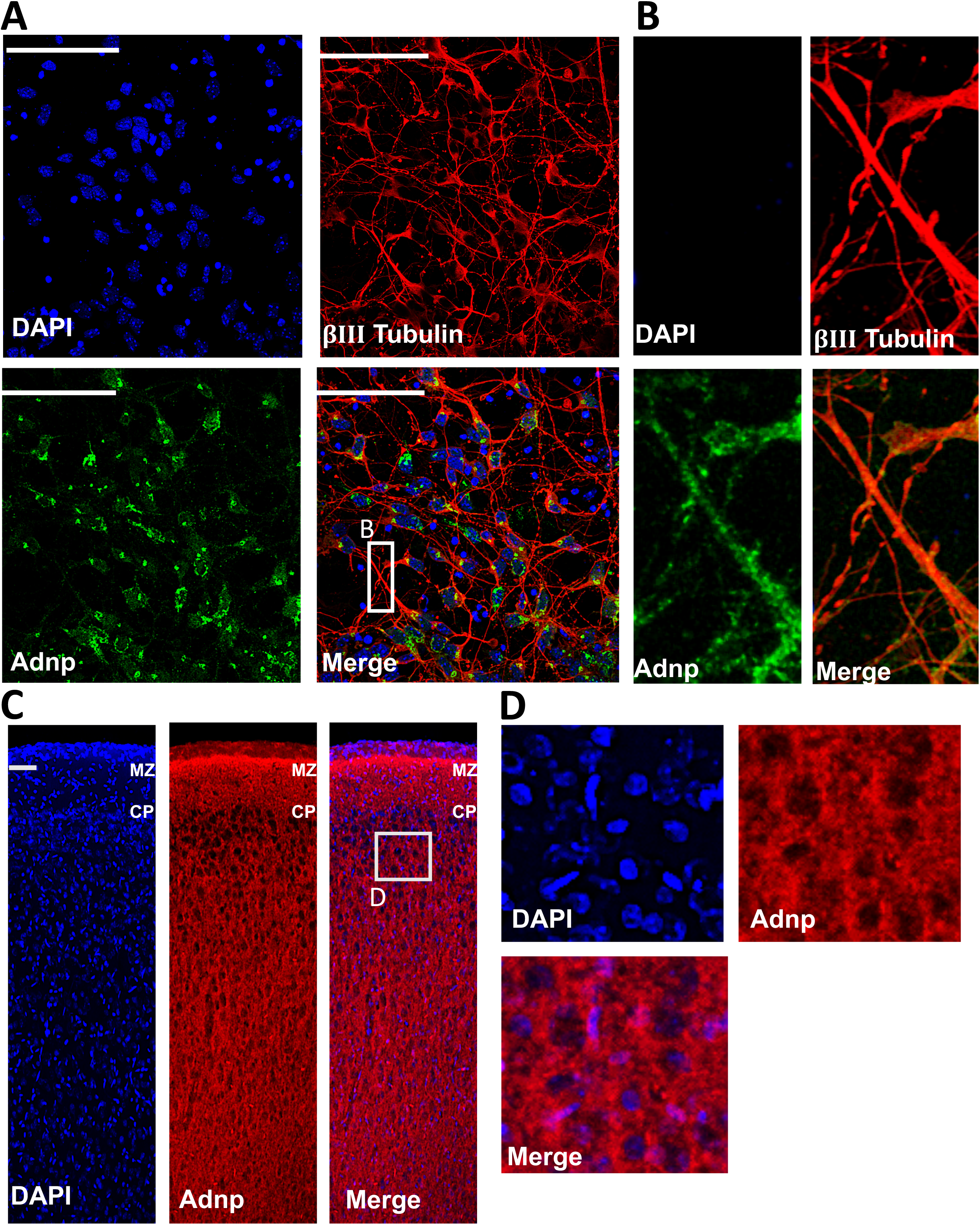
Immunofluorescence staining shows Adnp is expressed in primary cortical neurons and throughout the cortical plate. A) Primary cortical neurons harvested from E15.5 mouse embryos and cultured for 48 hours express Adnp in the cell soma and extending neurites. DAPI marks the nucleus, and βIII-tubulin is used as a cytoplasmic and neurite marker as well as to identify neurons. Scale bar= 100 µm B) High magnification of neurites expressing Adnp. C) Adnp is expressed throughout the cortical plate. The brain was harvested from a P15 mouse. Scale bar= 50 µm D) High magnification of cells in the cortical plate expressing Adnp. (3 independent staining experiments from 3 different litters)

In primary cortical neurons, we found that Adnp knockdown leads to a significant increase in length of the longest neurite, the neurite most likely to become the axon (Fig. 2A-B) (46), and no significant effect on length of the remaining neurites (Supplemental Fig. 2). Furthermore, we found a significant increase in neurite number in Adnp knockdown neurons (Fig. 2A and C). These results suggest that Adnp negatively regulates neurite initiation in the neurites likely to become dendrites, and negatively regulates neurite elongation in the neurite likely to become the axon. Defects were rescued by restoring Adnp expression in Adnp deficient cells (Fig. 2A-C). Rescue experiments were conducted using an shRNA resistant Adnp expression vector (“Adnp OE”) in combination with Adnp shRNA to restore Adnp expression. The control for our rescue was co-transfection of the resistant backbone vector (“control OE”) with scramble shRNA. These vectors were validated using western blot (Supplemental Fig. 1E). The rescue cells did not significantly differ from either the rescue control or shRNA scramble on measures of neurite number and length of the longest neurite, confirming that these defects were from loss of Adnp and not from off-target effects of the shRNA (Fig. 2A-C). Sholl analysis of neurite branching corroborated our neurite formation analysis, with increased intersections proximally in Adnp deficient neurons, indicating increased neurite initiation, and increased intersections distally by an average of 1 intersection, indicating increased elongation of a single neurite (Fig. 2D).

**Figure 2.**
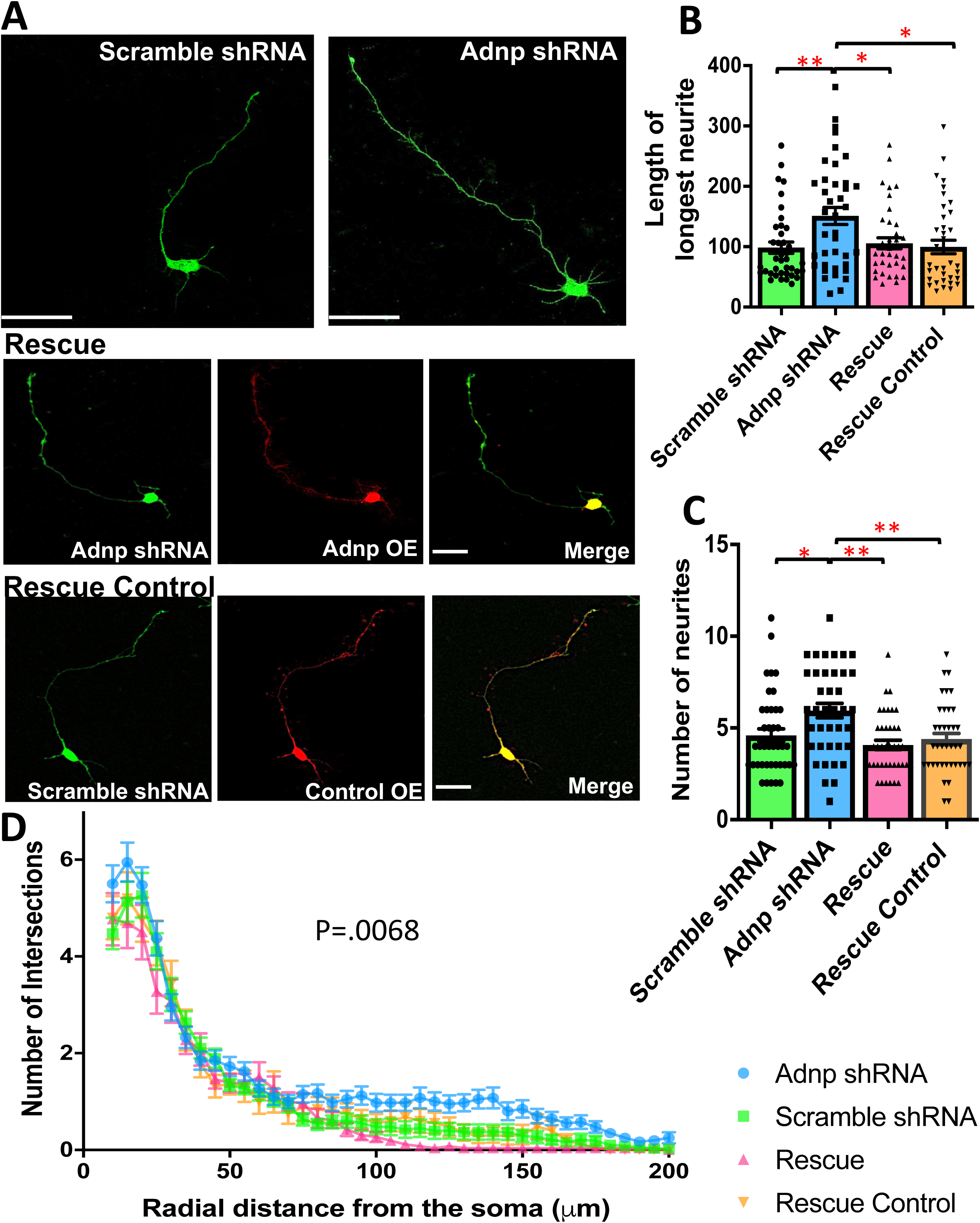
Adnp knockdown produces neurite initiation and elongation defects *in vitro*. A) Representative neurons expressing Scramble-shRNA-Venus, Adnp-shRNA-Venus, or the rescue groups. The rescue group co-expresses Adnp-shRNA-Venus and an Adnp-mScarlet expression vector (“Adnp OE”), which restores Adnp expression in Adnp deficient neurons; and the rescue control group which co-expresses Scramble-shRNA-Venus and the empty mScarlet backbone Adnp expression vector (“Control OE”). B) Quantification of length of the longest neurite. A one-way ANOVA showed a significant difference in length of the longest neurite between groups, F(3, 151)=4.904, p=0.003. Bonferroni post hoc comparison showed that Adnp shRNA neurons had significantly longer neurites (150.979*μ*m ± 14.233*μ*m) compared to Scramble shRNA (98.459*μ*m ± 9.305*μ*m, p=0.007), rescue (105.566*μ*m ± 9.258*μ*m, p=0.028), and rescue control neurons (101.316*μ*m ± 11.307*μ*m, p=0.013). Scramble shRNA neurons did not significantly differ from the rescue neurons (p=1.000) or the rescue control neurons (p=1.000). C) Quantification of neurite number. A one-way ANOVA showed a significant difference in number of neurites between groups, F(3, 152)=6.283, p<0.0001. Bonferroni post hoc comparison showed that Adnp shRNA neurons had significantly more neurites (5.949 ± 0.389) compared to Scramble shRNA (4.539 ± 0.354, p=0.018), rescue (4.075 ± 0.259, p=0.001), and rescue control neurons (4.395 ± 0.312, p=0.007). Scramble shRNA neurons did not significantly differ from the rescue neurons (p=1.000) or the rescue control neurons (p=1.000). D) Sholl analysis for neurite branching corroborates morphology analysis, with Adnp shRNA neurons having greater branching towards the soma, indicating increased neurite number, and a single branch point extending towards a greater distal length, indicating increased length of the longest neurite. A two-way repeated measures ANOVA revealed a significant difference between groups on neurite branching F(3, 125) = 4.245, p=0.0068. (n=40 measurements per group from 40 distinct neurons, 3 independent experiments per condition from 3 different litters). Scale bars = 50*μ*m.

### Adnp knockdown disrupts cortical neuritogenesis in vivo

We confirmed these results *in vivo* using IUE to knock down Adnp at E15.5 in neurons that migrate and mature into layer 2/3 pyramidal neurons in the somatosensory cortex (Supplemental Fig. 3) (47). We analyzed dendritic morphology including apical and basal dendrite length and number at P15, after neuritogenesis has proceeded to completion, roughly equivalent to our *in vitro* analysis (Fig. 3). These results were consistent with our *in vitro* analysis in that there was an increase in basal dendritic number on Adnp deficient neurons (Fig 3A-B), whereas *in vitro* there was an increase of neurite number (Fig. 2C). However, there was an additional deficit noted *in vivo* of decreased basal dendrite length on Adnp deficient neurons (Fig 3A and C). Both of these defects were rescued by restoring Adnp expression (Fig. 3A-C). Apical dendrite length was not affected (Supplemental Fig. 4).

**Figure 3.**
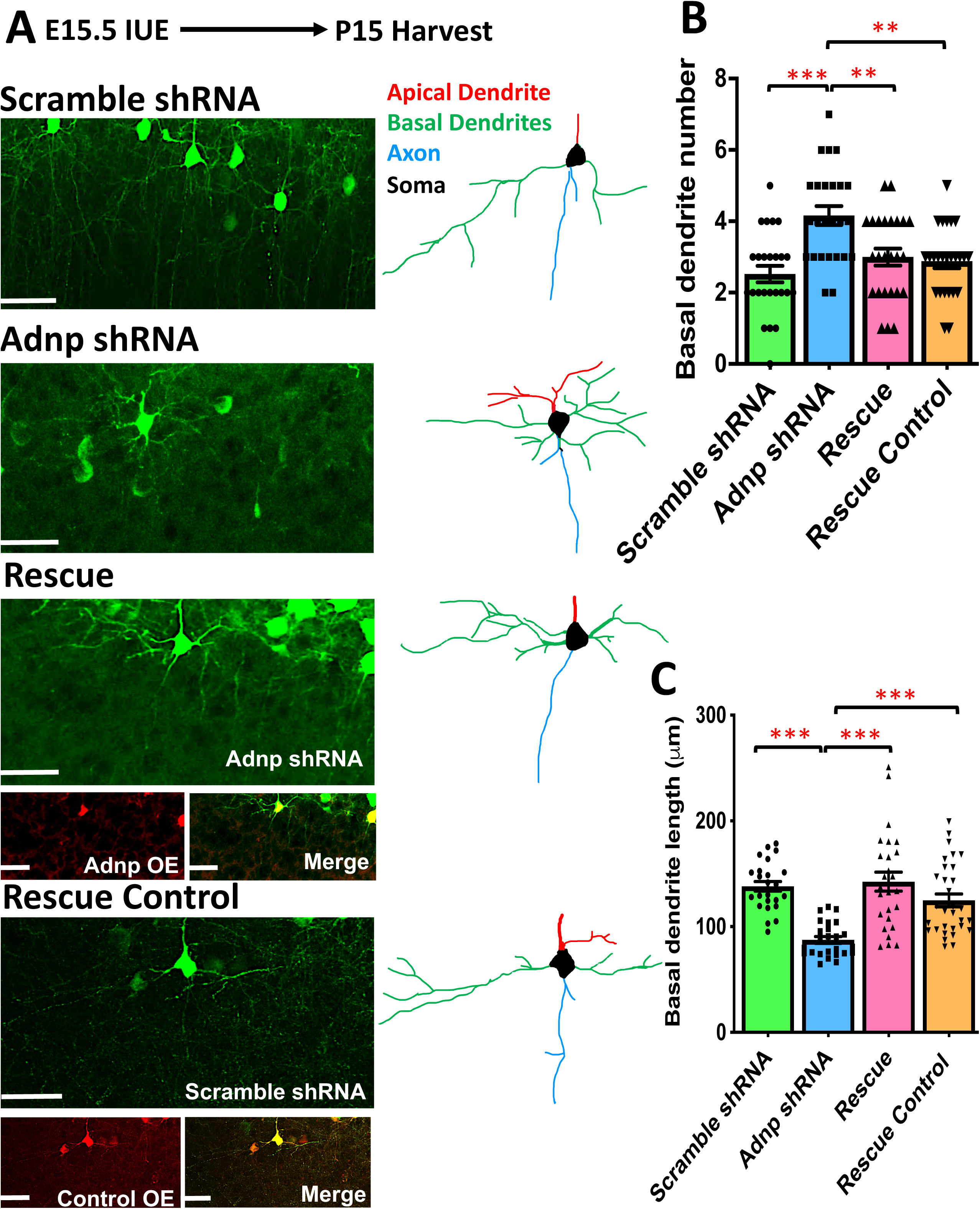
Adnp knockdown produces defects in basal dendrite development *in vivo*. A) Representative photos of layer 2/3 pyramidal neurons at P15 expressing Scramble shRNA, Adnp shRNA, Rescue, or Rescue control plasmids. Accompanying each representative photo is a color-coded tracing for clear visualization and comparison of dendritic morphology. Red= apical dendrite, green= basal dendrites, blue= axon, black= soma. B) Quantification of basal dendrite number between groups (n=25 neurons). A one-way ANOVA revealed a significant difference in basal dendrite number between groups, F(3, 96) = 9.287, p<0.0001. Bonferroni post hoc comparison showed that Adnp shRNA neurons had significantly more basal dendrites (4.160 ± 0.264) compared to Scramble shRNA (2.520 ± 0.232, p<0.001), rescue (3.000 ± 0.238, p=0.004), and rescue control neurons (2.880 ± 0.194, p=0.001). Scramble shRNA neurons did not significantly differ from the rescue neurons (p=0.890) or the rescue control neurons (p=1.000). Scale bars= 100*μ*m. IUE was performed at E15.5. C) Quantification of basal dendrite length between groups. A one-way ANOVA revealed a significant difference in basal dendrite length between groups, F(3, 103) = 15.376, p<0.0001. Bonferroni post hoc comparison showed that Adnp shRNA neurons had significantly shorter basal dendrites (87.478*μ*m ± 3.228*μ*m, n=25) compared to Scramble (138.064*μ*m ± 4.566*μ*m, p<0.0001, n=25), rescue (142.510*μ*m ± 8.982*μ*m, p<0.0001, n=26), and rescue control neurons (124.858*μ*m ± 6.067*μ*m, p<0.0001, n=31). Scramble shRNA neurons did not significantly differ from the rescue neurons (p=1.000) or the rescue control neurons (p=0.766). Measurements for basal dendrite length were taken from 15 distinct neurons. (3 independent experiments per condition from 3 different litters).

Next, we assessed axon length at P15, after axons have crossed the midline and traveled to their final destinations and integrated into their appropriate cortical circuits. We cut coronal brain slices of the electroporated area and traced the axon bundle from the midline to its termination, which was defined as no measurable Venus fluorescence. There was no difference in density of axons crossing the midline as measured by width of the midline bundle in Adnp shRNA compared to control (Fig. 4A-C). This suggests that there is no difference in number of neurons sending their axons to the contralateral hemisphere. We found an increase in length of the axon bundle after crossing the midline in Adnp deficient neurons, again confirming the phenotype observed *in vitro* (Fig. 4A, B, D-F). We also observed increased innervation throughout the entire opposing cortex from axons deficient in Adnp compared to control (Fig. 4G-H).

**Figure 4.**
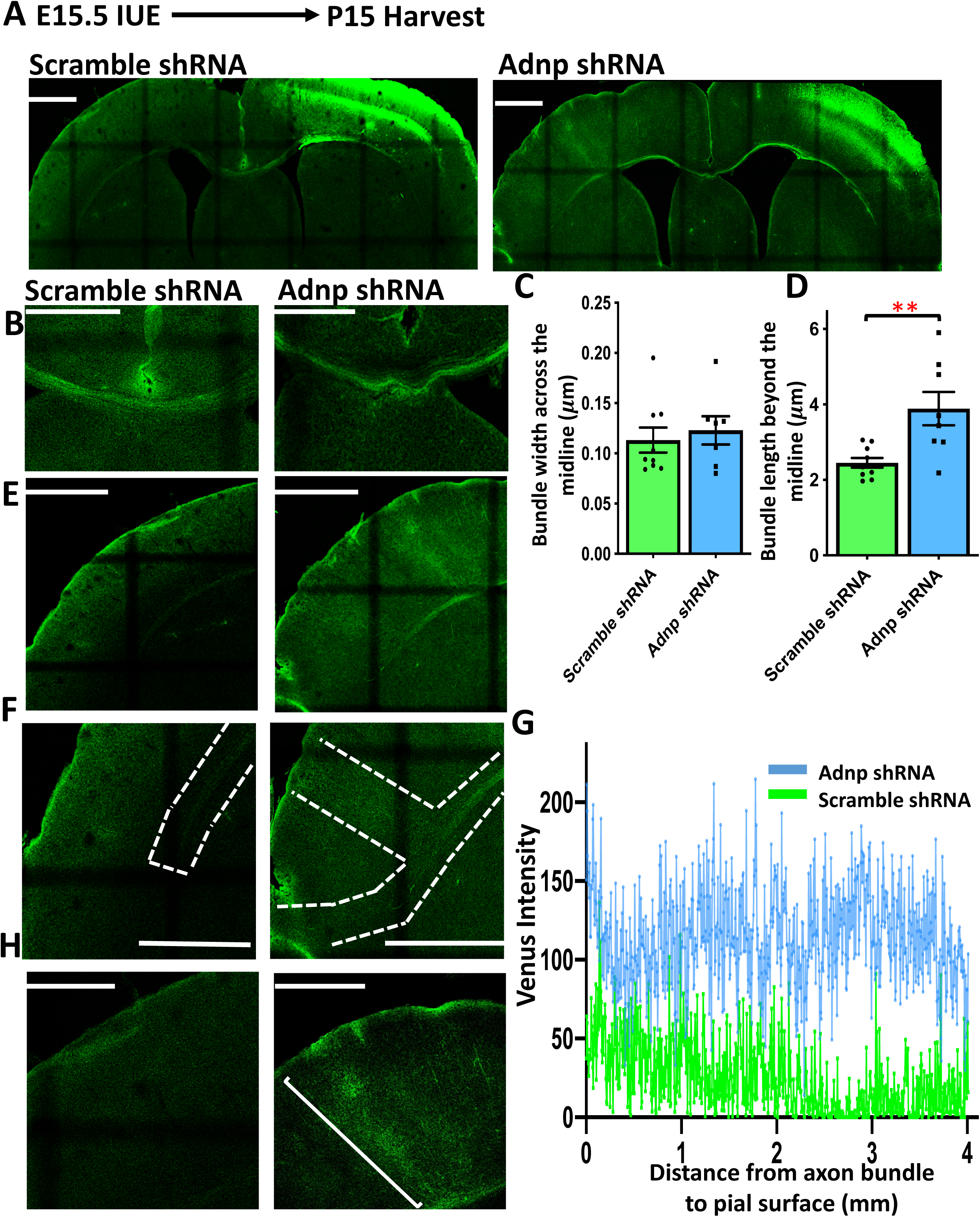
Adnp knockdown increases interhemispheric cortical axon length and innervation throughout the opposing cortex. IUE was performed at E15.5. and brains were collected at P15. A) Low magnification representative photos of P15 brain slices from Scramble shRNA and Adnp shRNA expressing mice. All high magnification photos were taken of these slices. B) High magnification representative photos of the axon bundle crossing the midline. C) Quantification of axon bundle width as it crosses the midline. An independent samples t-test showed that there was no significant difference between bundle width between Scramble shRNA (0.113mm ± 0.012mm, n=9) and Adnp shRNA (0.122mm ± 0.014mm, n=7) expressing axons (95% CI, −0.050 to 0.031), t(14)= −0.516, p=0.614, indicating that there is no difference in number of axons crossing the midline. D) Quantification of length of the axon bundle after crossing the midline. An independent samples t-test showed that Adnp shRNA expressing neurons extended axons a significantly greater distance across the midline (3.887mm ± 0.441mm, n=8) compared to Scramble shRNA expressing neurons (2.453mm ± 0.128mm, n=10) (95% CI, −2.318 to −0.550), t(16)= −3.438, p=.0003. E) High magnification representative photos of the opposing hemispheres from the ventricle. F) High magnification representative photos of the area where the axon bundles terminate. G) Quantification of axon innervation throughout the opposing cortical hemisphere as measured by Venus intensity. H) High magnification representative photos of the opposing hemisphere from the axon bundle. Scale bars= 1 mm. (reported n refers to number of brain slices, 3 independent experiments per condition from 3 different litters).

### Adnp knockdown in vivo disrupts apical dendrite orientation

At early postnatal stages, cortical neurons have a relatively immature morphology characterized by a single apical dendrite extended at roughly a 90° angle with respect to the cortical plate (Fig. 5A). The angle of the apical dendrite at this stage influences both neuronal function and synaptic connectivity as neural networks and initial synaptic contacts are forming (48). Analysis of Adnp deficiency *in vivo* by using IUE at E15.5 and analysis at P3 revealed a defect in the angle of the apical dendrite. Scramble shRNA cells extended their apical dendrites straight from the soma to form a roughly 90° angle with the cortical plate, with a tight distribution of angles spanning from less than −60° to 60° (Fig. 5B). This deviation significantly differed from Adnp deficient neurons which extended apical dendrites at many different angles, often with sharp bends that fell across a broad distribution from −30° to 30° (Fig. 5C). The angle of the apical dendrite was restored by rescue of Adnp expression using our validated vectors, confirming that Adnp signaling is responsible for this defect (Fig. 5D-E). The apical dendrite angle distribution for rescue and rescue control cells also fell between −60° and 60°, the same as shRNA scramble cells. We calculated the average deviation from the expected 90° across groups and found that Adnp deficient neurons indeed had a significantly greater deviation compared to Scramble shRNA, rescue, and rescue control groups (Fig. 5F). Furthermore, we found that apical dendrites present on knockdown neurons were significantly wider with respect to the soma compared to controls (Fig. 5G). These results further reveal Adnp’s complex involvement in establishing neuronal morphology and potentially network connectivity. We found that these deficits are sustained throughout P15 and were also rescued due to restoration of Adnp expression in Adnp deficient cells (Supplemental Fig. 5).

**Figure 5.**
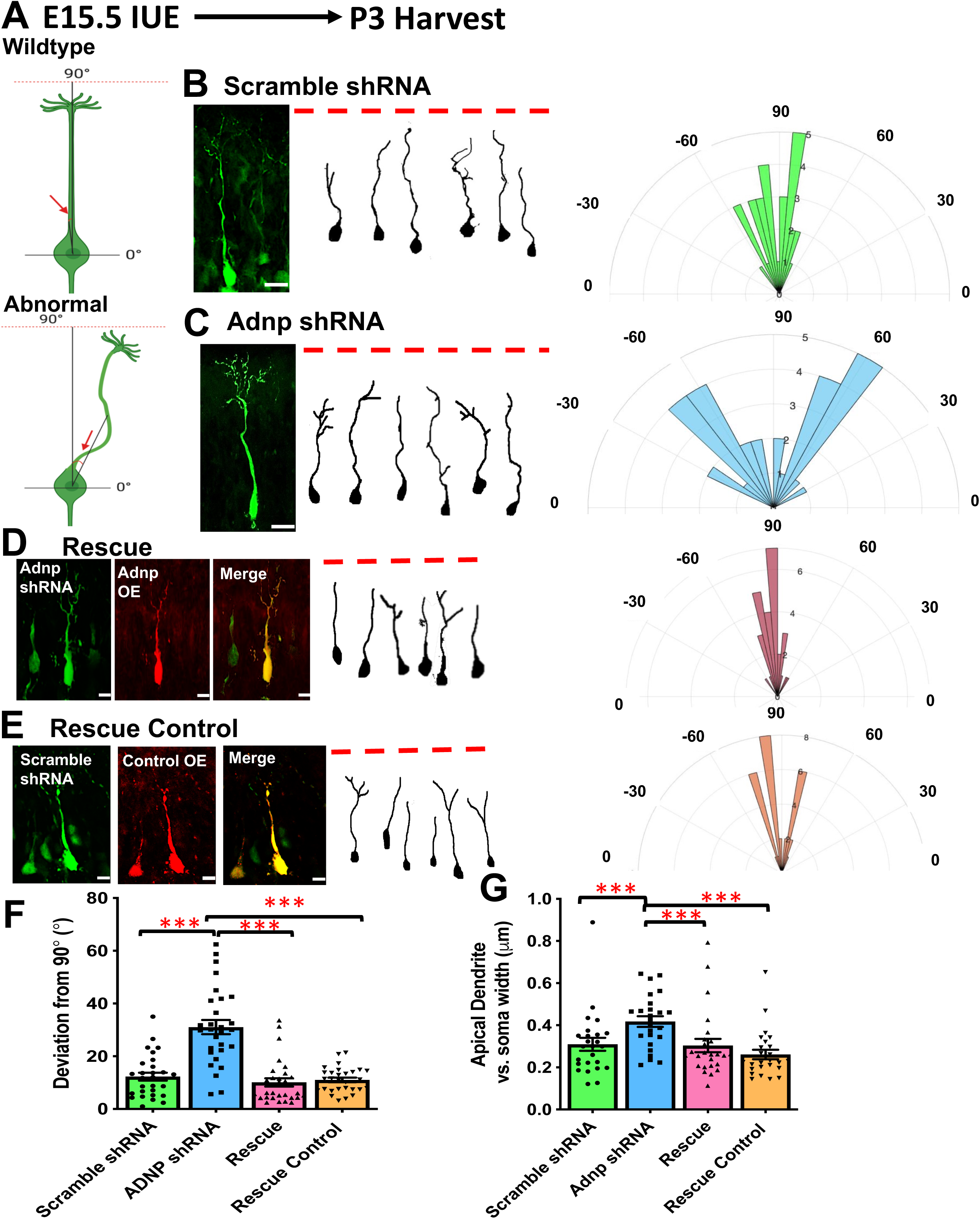
Adnp knockdown disrupts apical dendrite development *in vivo*. IUE was performed at E15.5 and brains were collected at P3. A) Schematic for how the angle of the apical dendrite was measured. Angles were measured by drawing one line from the soma perpendicular to the cortical plate and another line from the soma to the middle of the apical dendrite. The angle between these lines was measured. The red dotted line represents the pial surface. Representative photos of layer 2/3 pyramidal neurons expressing Scramble shRNA (B), Adnp shRNA (C), rescue plasmids Adnp-shRNA-Venus and Adnp-mScarlet (“Adnp OE”) (D), and rescue control plasmids Scramble-shRNA-Venus and mScarlet backbone vector (“Control OE”) (E). All photos are accompanied by tracings of 6 representative neurons for clear visualization of apical dendrite morphology, and by polar histograms which quantify the angles at which the apical dendrites extended with respect to the cortical plate. F) Quantification of the angle of the apical dendrite from the soma. These graphs depict the degrees by which the apical dendrites deviate from the expected 90°, with the greater angle indicating a greater defect. A one-way ANOVA showed that there was a difference in apical dendrite angle deviation between groups F(3,109)= 26.751, p<0.0001. Bonferroni post hoc comparison revealed that Adnp shRNA expressing neurons had a significantly greater deviation from 90° (27.807° ± 2.244°, n=26) compared to Scramble shRNA (12.287° ± 1.489°, p<0.0001, n=29), rescue (10.070° ± 1.554°, p<0.0001, n=29), and rescue control (11.015° ± 0.881°, p<0.0001, n=29). Scramble shRNA neurons did not significantly differ from the rescue neurons (p=1.000) or the rescue control neurons (p=1.000). G) Quantification of the width of the apical dendrites with respect to the soma, which was calculated by diving the apical dendrite width by the soma width to account for soma width variation. A one-way ANOVA showed that there was a significant difference between groups, F(3,88) = 20.374, p<0.0001. Bonferroni post hoc comparison showed that Adnp shRNA neurons had significantly wider apical dendrites with respect to the soma (0.418*μ*m ± 0.025*μ*m, n=25) compared to Scramble shRNA (0.277*μ*m ± 0.019*μ*m, p<.0001, n=23), rescue (0.253*μ*m ± 0.015*μ*m, p<.0001, n=24), and rescue control neurons (0.229*μ*m ± 0.013*μ*m, p<0.0001, n=24). Scramble shRNA neurons did not significantly differ from the rescue neurons (p=1.000) or the rescue control neurons (p=0.525). Scale bars= 10 *μ*m. (reported n refers to both number of apical dendrites and number of distinct neurons, 3 independent experiments per condition from 3 different litters).

### Ex vivo time-lapse live imaging reveals disruption of neuritogenesis dynamics in Adnp deficient neurons

To pinpoint when the primary morphological deficits develop for Adnp deficient neurons, as defects at one stage effect all other stages, we observed cortical sections as neurons completed neurogenesis at E17.5 (Supplementary Fig. 6A) and were migrating at E18.5 (Supplementary Fig. 6B) and we found no differences in cortical positioning. At our apical dendrite analysis at P3, all neurons had arrived at their correct positions in the cortical plate, suggesting no defects in neuronal migration (Supplementary Fig. 6C). Defects occurring at the onset of neuritogenesis, at P0, likely underlie our observed later deficits at P3 and P15 in axo- and dendritogenesis. To investigate the highly dynamic process of neuritogenesis in more detail, we performed *ex vivo* time-lapse live imaging on living brain slices harvested from P0 cortices of mice that had undergone IUE at E15.5 to introduce either Adnp shRNA or scramble shRNA. At P0, layer 2/3 neurons have terminated migration and settled in the cortical plate where they are just beginning neurite initiation. Time-lapse live imaging allowed us to detect many interesting flaws in Adnp deficient neurons compared to control as neuritogenesis occurred that the use of fixed samples did not (Fig 6, Supplemental movies 1 and 2). Scramble shRNA neurons were highly dynamic, displaying rapid changes in morphology with extension and retraction of filopodia and lamellipodia which were eventually stabilized and then elongated (Fig. 6A-B, Supplemental Video 1). Adnp shRNA neurons had on average one primary neurite at the beginning of imaging that was already significantly longer than the primary neurites present on control cells (Fig. 6A-B, Supplemental Video 2). As imaging progressed, instead of the rapid formation and retraction of primitive neurites, the primary neurite on Adnp deficient neurons either remained stable or slowly grew with reduced maximum growth and retraction velocity (Fig 6C-E), resulting in a significantly shorter length of growth throughout the video compared to scramble shRNA (Fig. 6F). This slow yet consistent growth of a single neurite, as opposed to rapid shrinkage and growth seen in controls, likely explains the eventual increase in axon length *in vitro* and *in vivo* Adnp deficient neurons. These results suggest that Adnp shRNA neurons have defects in the dynamics necessary to produce appropriate neurite elongation.

**Figure 6.**
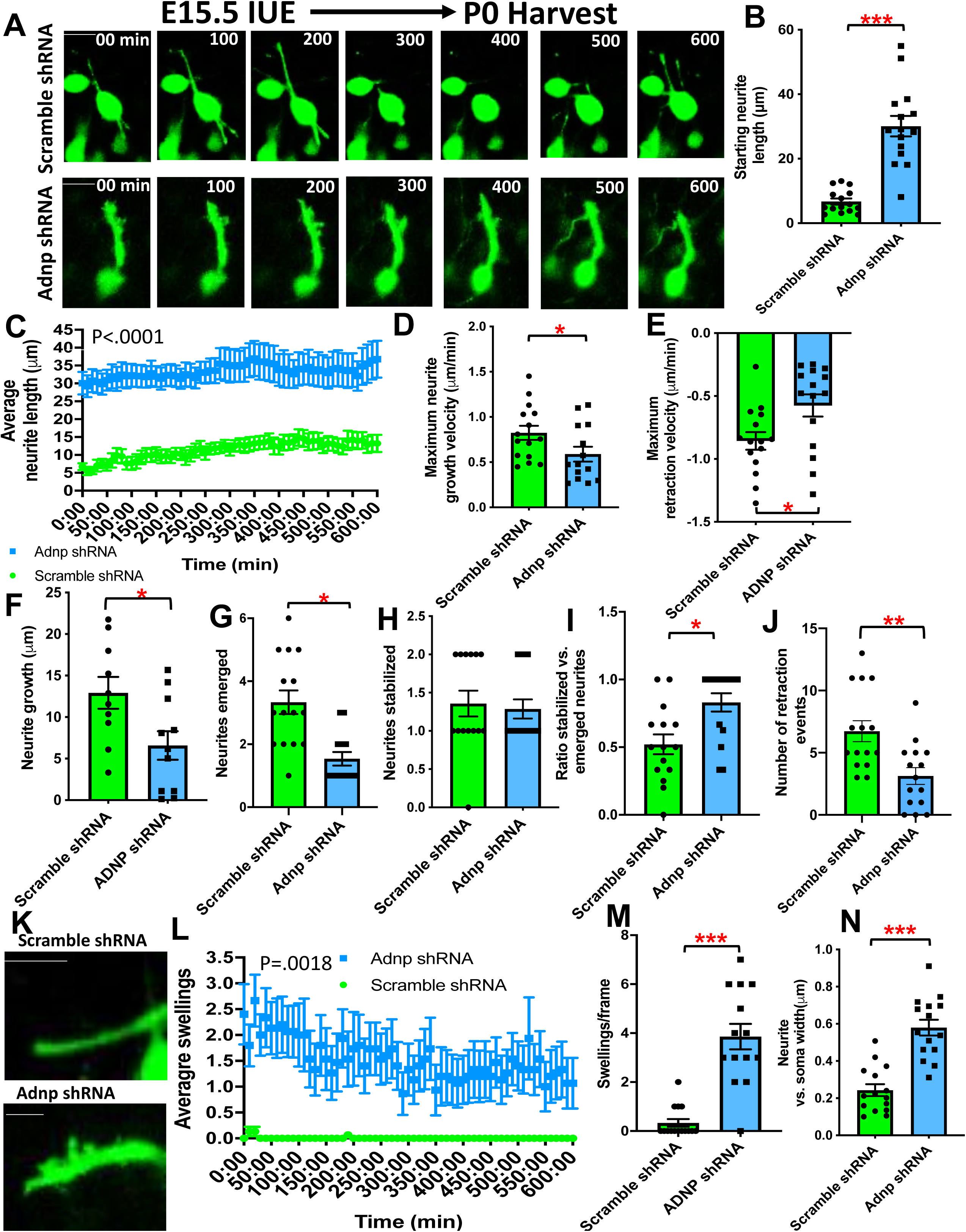
Live imaging reveals that deficits in neuritogenesis due to Adnp knockdown begin at P0. IUE was performed at E15.5 and brains were collected at P0. A two-tailed independent samples t-test was used for all statistical analyses unless otherwise indicated. (reported n refers to number of distinct neurons, 3 independent experiments per condition from 3 different litters). A) Representative time-lapse images of Scramble shRNA and Adnp shRNA expressing neurons over the course of 600 minutes, or 10 hours. Each panel is time-stamped with the minutes. Scale bar = 10*μ*m. B) Quantification of starting length of the neurites present at the start of imaging. Adnp shRNA expressing neurons had significantly longer neurites (30.090*μ*m ± 3.160*μ*m, n=10) compared to Scramble shRNA expressing neurons (6.723*μ*m ± 0.938*μ*m, n=10), (95% CI 16.62 to 30.12), t(28) = 7.090, p<0.0001. C) Quantification of neurite length throughout the 10-hour imaging. A two-way repeated measures ANOVA revealed a significant difference between Adnp shRNA and Scramble shRNA neurite growth, F(1, 28) = 29.01, p<0.0001, n=10. D) Quantification of the maximum growth velocity (Vmaxg) of neurites. Adnp shRNA expressing neurons had a significantly slower Vmaxg (0.589*μ*m/min ± 0.083*μ*m/min, n=14) compared to Scramble shRNA expressing neurons (0.824*μ*m/min ± 0.079*μ*m/min, n=15), (95% CI 0.002 to 0.470), t(27) = 2.067, p=0.048. E) Quantification of the maximum retraction velocity (Vmaxr) of neurites. Adnp shRNA expressing neurons had a significantly slower Vmaxr (−0.575*μ*m/min ± 0.088*μ*m/min) compared to Scramble shRNA expressing neurons (−0.857*μ*m/min ± 0.070*μ*m/min), (95% CI −0.512 to - 0.052), t(28) = −2.511, p=0.018, n=15. F) Quantification of neurite growth throughout imaging. Adnp shRNA expressing neurons had significantly less total growth (6.585*μ*m ± 1.713*μ*m, n=11) compared to Scramble shRNA expressing neurons (12.934*μ*m ± 1.918*μ*m, n=10), (95% CI 0.984 to 11.713), t(19) = 2.477, p=0.023. G) Quantification of number of neurites emerged throughout the imaging. Adnp shRNA expressing neurons had significantly fewer neurites emerge (1.857 ± .376, n=14) compared to Scramble shRNA (3.333 ± 0.374, n=15), (95% CI 0.387 to 2.565), t(27) = 2.782, p=0.010. H) Quantification of neurites stabilized (present and elongating) and the conclusion of imaging. There was no significant difference between Adnp shRNA neurons (1.286 ± 0.125) and Scramble shRNA neurons (1.357 ± 0.169), (95% CI −0.361 to 0.504), t(26) = 0.339, p=0.737, n=14. I) Quantification of the ratio of stabilized to emerged neurites. This ratio indicates the likelihood of each neurite that emerges being stabilized. Adnp shRNA neurons had a significantly higher rate of stabilization (0.831 ± 0.068) compared to Scramble shRNA (0.521 ± 0.285), (95% CI −0.514 to −0.105), t(28) = −3.098, p =0.004, n=15. J) Quantification of number of retraction events throughout the imaging. Adnp shRNA expressing neurons retracted significantly fewer neurites (3.133 ± 0.675) compared to Scramble shRNA (6.733 ± 0.842), (95% CI 1.389 to 5.811), t(28) = 3.335, p=0.002, n=15. K) Representative photos of swellings on the apical dendrites. Scale bar= 5*μ*m. L) Quantification of swelling events on the primary neurite throughout imaging. A two-way repeated measures ANOVA revealed a statistically significant difference between groups, F(1, 28)=11.95, p = 0.0018. M) Quantification of amount of swellings per frame of imaging (10 minutes). Adnp shRNA neurons had significantly more swellings (3.857 ± 0.523, n=14) compared to Scramble shRNA (0.333 ± 0.159, n=15), (95% CI −4.613 to −2.435), t(27) = - 6.641, p<0.0001. N) Quantification of primary neurite width. Adnp shRNA neurons had significantly wider primary neurites (4.300*μ*m ± 0.374) compared to Scramble shRNA (2.728*μ*m ± 0.216), (95% CI −2.357 to −0.587), t(28) = −3.405, p=0.002, n=15.

Scramble shRNA cells on average had significantly more neurites emerge throughout the 10 hour time span than Adnp shRNA cells (Fig. 6G) yet the final number of neurites stabilized was not significantly different (Fig. 6H), resulting in a ratio of emergence to stabilization being significantly higher in Adnp shRNA neurons compared to scramble shRNA neurons (Fig. 6I). These results suggest that Adnp shRNA neurons have issues producing the necessary dynamics of neurite initiation. Furthermore, Adnp deficient neurons also had a significantly lower neurite retraction frequency (Fig. 6J). The higher ratio of neurite stabilization and the decreased retraction frequency suggests that in Adnp deficient cells, each neurite that emerges is more likely to be stabilized, regardless of the intrinsic properties of that neurite that might not make it a good candidate for stabilization. This explains the eventual increase in neurite number seen *in vitro* and *in vivo* in Adnp deficient neurons. Taken together, these results suggest serious flaws in the necessary dynamics that drive both proper neurite initiation and elongation.

A final interesting morphological phenotype revealed by live imaging was neuritic swelling and appearance of dilations along the length of neurites present on Adnp shRNA neurons, as well as the width of the primary neurites present on Adnp shRNA neurons (Fig. 6K-N). These swellings rarely appeared in shRNA scramble neurons. Adnp deficient neurons had significantly more swellings per frame (Fig. 6L-M), which were defined as outgrowths from the primary neurite measuring more than one micron that did not mature into branching points, than scramble shRNA cells. Primary neurites on Adnp deficient cells were also significantly wider compared to controls (Fig. 6N). This increase in apical dendrite width was sustained at P3 and P15 where it was able to be rescued by restoring Adnp expression in Adnp deficient cells (Fig 5G, Supplemental Fig. 5I). These results suggest intracellular issues within Adnp deficient neurons, perhaps relating to transport of organelles large enough to produce these swellings, such as mitochondria and Golgi.

### Adnp knockdown disrupts properties of dendritic spines in vivo

Next, to assess later stages of neuronal morphogenesis and maturation, we analyzed dendritic spine density and morphology at P15. A previous study showed a slight, yet significant decrease in dendritic spine density in the Adnp haploinsufficient mouse (34). We further assessed dendritic spine morphology using a geometric quantification system (Fig. 7A) (42). We found that Adnp deficient neurons had a significant decrease in proportion of mushroom type spines, which are generally considered the most mature and the site of functional synaptic contacts (16), and an increase of other spine morphologies such as thin (Fig 7B-D). We also confirmed a decrease in dendritic spine density in our model (Fig. 7E) consistent with the Adnp haploinsufficient mouse (34). Our results suggest several dendritic spine properties are altered due to loss of Adnp, potentially effecting synaptic efficiency.

**Figure 7.**
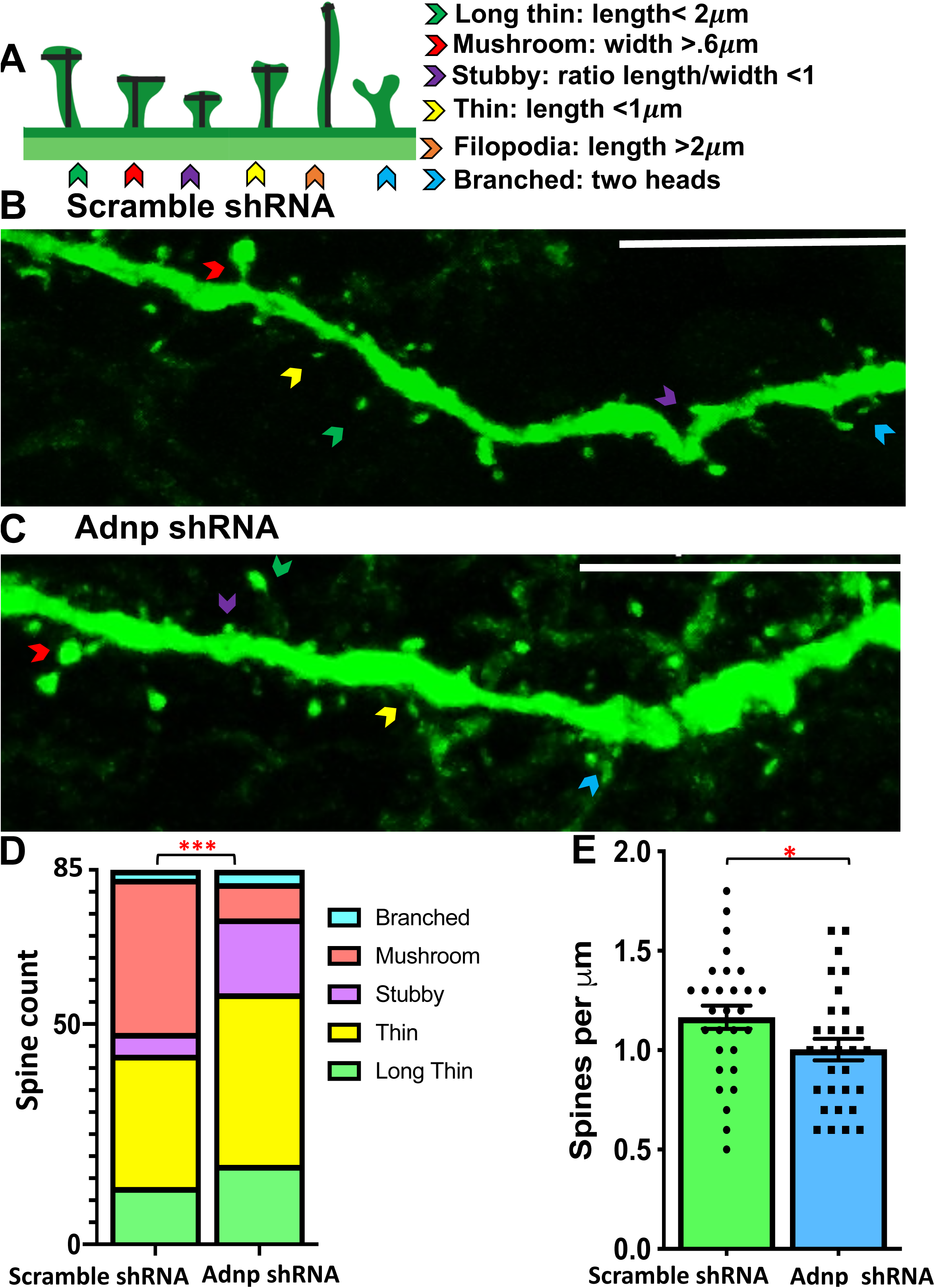
Adnp knockdown disrupts dendritic spine development *in vivo*. IUE was performed at E15.5 and brains were collected at P15. Scale bars= 10*μ*m A) Schematic depicting quantitative criteria for spine morphology categories, modified from Rischer et al. (42). B-C) Representative photo of dendritic spines present on a neuron expressing Scramble shRNA (B) or Adnp shRNA (C). D) Quantification of proportion of spines with differing morphologies. The bar graph displays the spine count per morphology as a proportion of total spines. A chi-squared test for homogeneity revealed there was significant difference in proportions of spine morphologies between Scramble shRNA and Adnp shRNA expressing neurons (p<0.001). The percentages of spine morphologies were as follows: long thin (Scramble shRNA 15.7%, Adnp shRNA 22%), thin (Scramble shRNA 36.1%, Adnp shRNA 47.6%), stubby (Scramble shRNA 6.0%, Adnp shRNA 20.7%), mushroom (Scramble shRNA 42.2%, Adnp shRNA 9.8%). There were only two cases of branched spines on Scramble shRNA dendrites and 3 on Adnp shRNA dendrites, and therefore this morphology was excluded from statistical analysis due to n<5. E) Quantification of dendritic spine density per *μ*m. An independent samples t-test revealed a slight yet significant decrease in dendritic spine density in Adnp shRNA expressing neurons (1.003± 0.05456) compared to Scramble shRNA (1.166 ± 0.05794), (95% CI −0.3214 to −0.002944), t(57)= 2.039, p= 0.0460. (n=85 dendritic spines from 25 dendrites from 10 neurons per group, 3 independent experiments per condition from 3 different litters).

### Adnp deficient pyramidal neurons show increased spontaneous calcium signaling

Although our dendritic spine analysis suggests that there may be less synaptically functional spines on Adnp deficient neurons; the increased axon length, innervation to opposing cortical layers, and basal dendrite numbers in Adnp deficient neurons may actually provide more surface area for these neurons to form synaptic contacts, negating the effect of the dendritic spines. To directly test how loss of Adnp affects neuronal function we performed *ex vivo* calcium imaging. Mice underwent IUE at E15.5 to introduce either Adnp or scramble shRNA tagged with an mScarlet reporter in combination with the genetically encoded calcium indicator GCaMP6s. Brains were harvested for imaging experiments at 2 months old and spontaneous calcium activity was measured (Fig 8A). Circular regions of interest were placed over the cell somas, and average fluorescence values were extracted at each 30ms time frame. We found that Adnp deficient neurons had significantly greater calcium influx, as measured by percent fluorescence change from baseline, compared to control (Fig. 8A-B). We also found that a greater percentage of Adnp deficient neurons had influxes of calcium large enough to indicate an action potential compared to control, although this change was not statistically significant (Fig. 8B-C) (44). These results suggest that although Adnp deficient neurons have altered spine properties, they have increased spontaneous calcium signaling compared to controls.

**Figure 8.**
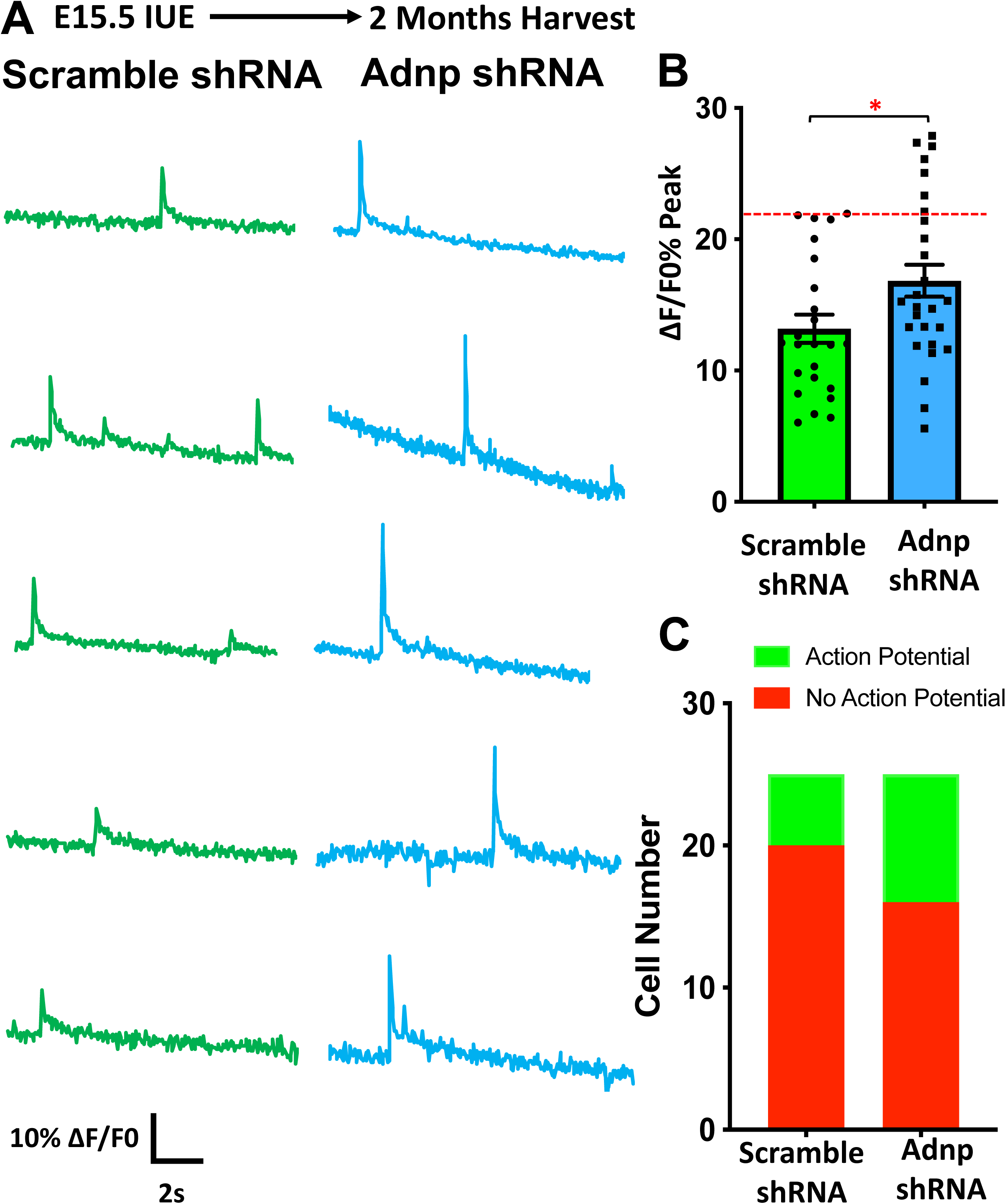
Adnp deficient cells exhibit increased spontaneous calcium signaling. IUE was performed at E15.5 and imaging took place at P30. A) Fluorescence percent change traces from 5 representative cells from Scramble or Adnp shRNA expressing neurons. B) Quantification of fluorescence percent change peaks. The red dotted line indicates the peak fluorescence threshold to reliably predict single action potential firing as defined by Chen et al. as 23% ± 3.2% for GCaMPs (44). An independent samples t-test revealed that Adnp shRNA neurons had a significantly higher peak fluorescence percent change (16.844% ± 1.217%, n=27) compared to Scramble shRNA (13.186% ± 1.071%, n=25), (95% CI −6.95237 to −0.36367), t(49)= −2.231, p= 0.030. C) Quantification of the proportions of cells that did or did not reach fluorescence threshold to indicate and action potential. 20% of Scramble shRNA expressing neurons (n=25) and 33.3% of Adnp shRNA neurons (n=27) reached fluorescence threshold, however this is not a statistically significant difference using a chi-squared test for homogeneity (p=0.279). (3 independent experiments per condition from 3 different litters).

### GRAPHIC reveals increased interhemispheric cortical connectivity and excitatory shaft synapses in Adnp deficient neurons

Increased axon innervation from pyramidal neurons to opposing cortical neuronal populations was suggested by our axon tracing experiments. Furthermore, our calcium imaging suggests hyperexcitability of layer 2/3 neurons. To test whether this apparent increase in neuronal excitability and axon innervation correlated to increased connectivity between layer 2/3 neurons, we utilized a novel tracing technique, “GPI anchored reconstitution-activated proteins highlight intercellular contacts” GRAPHIC, which delineates synaptic contacts between neurons using a GFP reconstitution method (49). GRAPHIC utilizes expression vectors encoding GPI-anchored membrane proteins that display complementary fragments of the GFP protein (49). Therefore, GFP is specifically reconstituted at the contact area between two cells expressing complementary plasmids, which has been validated as synapse-specific when transfected into neurons (49) (Fig. 9A). A pair of complementary GRAPHIC plasmids were injected into opposing cortical hemispheres (Fig. 9B), one with an mCherry reporter and one with an H2B-mCherry reporter, at E15.5 in combination with either Adnp or scramble shRNA containing a mTagBFP2 reporter. Brains were harvested and mCherry+, BFP+ and GFP puncta+ neurons were analyzed at P30 (Fig. 9C), when pyramidal cortical neurons are morphologically and synaptically mature (50). Adnp deficient dendrites had significantly more puncta compared to Scramble shRNA dendrites (Fig. 9C-D). We also observed a dramatic, significant increase in shaft puncta in Adnp deficient neurons compared to Scramble shRNA (Fig. 9E). In fact, the percentages of spine vs. shaft puncta were almost the complete opposite in Adnp deficient compared to scramble shRNA dendrites. Taken together, these results suggest Adnp deficient neurons have increased interhemispheric contacts from excitatory neurons, although the vast majority of these contacts are with the dendritic shaft, corroborating our calcium imaging.

**Figure 9.**
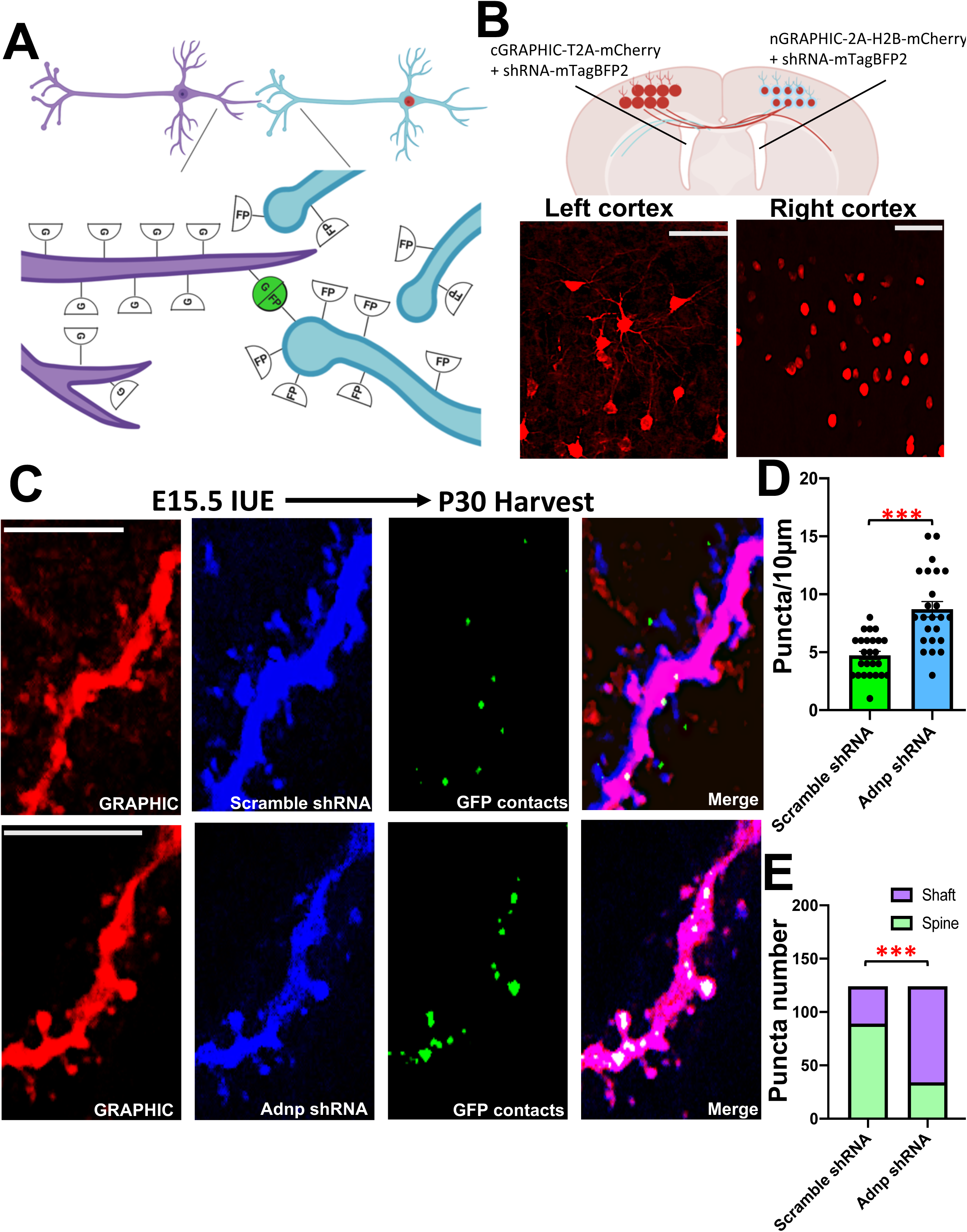
GRAPHIC reveals increased excitatory interhemispheric cortical connectivity and shaft synapses on Adnp deficient basal dendrites. A) Schematic for GRAPHIC’s GFP reconstitution system between neurons in the left and right cortical hemispheres. GFP puncta are exclusive to sites of synaptic contact between these two populations of neurons. Figure created using biorender.com. B) Schematic and validation of GRAPHIC transfection of layer 2/3 pyramidal neurons. The left hemisphere is mCherry positive and the right hemisphere is H2B-mCherry positive. Scale bar = 1 mm. Figure created using biorender.com. C) High magnification representative photos of basal dendrites with synaptic contacts. Scale bars = 10*μ*m. D) Quantification of GFP puncta per 10*μ*m. An independent samples t-test revealed that Adnp shRNA neurons (8.708 ± .669, n=24 measurements from 20 distinct dendrites from 12 different neurons) had significantly more GFP puncta compared to Scramble shRNA neurons (4.720 ± .349, n=25 measurements from 22 distinct dendrites from 19 different neurons), (95% CI 2.488 to 5.488), t(47)=5.349, p<0.0001. E) Quantification of the proportions of puncta on the dendritic shaft vs. spines. A chi-squared test for homogeneity revealed there was significant difference in proportions of shaft vs. spine synapses in Adnp shRNA (27.42% spine, 72.58% shaft) compared to Scramble shRNA neurons (71.77% spine, 28.23% shaft), p<0.0001. (2 independent experiments per conditions from 2 different litters).

### Adnp travels from the nucleus to the cytoplasm as primary cortical neurons undergo differentiation

To probe the mechanism for how Adnp may regulate neuritogenesis, we assessed its expression pattern as primary cortical neurons undergo differentiation and neurite formation. Adnp has differential functions, as a transcription factor in the nucleus and a cytoskeleton-interaction protein in the cytoplasm, and cell-type-specific expression patterns depending on the developmental stage of the cell (36, 37, 51–53). We were interested in assessing the expression pattern of Adnp in cortical neurons with a spherical, immature morphology vs. a mature, post-neurite formation morphology to provide information regarding which of Adnp’s roles, nuclear, cytoplasmic, or both are important for neuronal morphogenesis and maturation. To assess Adnp subcellular localization of immature neurons we dissected mouse cortices from E14.5 embryos and cultured them as neurospheres, allowing for the proliferation of neuronal stem cells. Neurospheres were kept intact and fixed after 14 days in culture, then we performed immunofluorescence staining (Fig. 10A). We compared neurospheres to primary cortical neurons harvested from E15.5 embryos plated on PDL and laminin-coated coverslips to encourage neuritogenesis. Primary cortical neurons were fixed after 48 hours, while in the late stages of neurite elongation, and immunofluorescently stained (Fig. 10B). We compared the staining patterns of Adnp in immature vs. mature neurons using three methods. Firstly, we compared the staining patterns using quantification of Adnp fluorescence intensity based on cellular compartment in neurospheres (Fig. 10C) and mature neurons (Fig. 10D). Secondly, we compared the fluorescence profiles of all fluorophores in a representative immature and mature neuron (Fig. 10E-F). Lastly, we compared the ratio of Adnp fluorescence intensity in the nucleus vs. the cytoplasm of neurospheres and mature neurons (Fig. 10G). These methods all show the same clear localization differences, that Adnp fluorescence is strongly distributed in the nucleus of immature neurons but mainly in the cytoplasm of neurons that have undergone neuritogenesis. In mature neurons, Adnp is most heavily localized to the perinuclear region and at the base of developing neurites (Fig. 10B). Adnp is still present in the nucleus of mature neurons, but it’s clear movement towards the cytoplasm as cortical neuronal maturation proceeds suggests Adnp may take on roles beyond that of a transcription factor, traveling to the cytoplasm during this specific time point to promote neuritogenesis.

**Figure 10.**
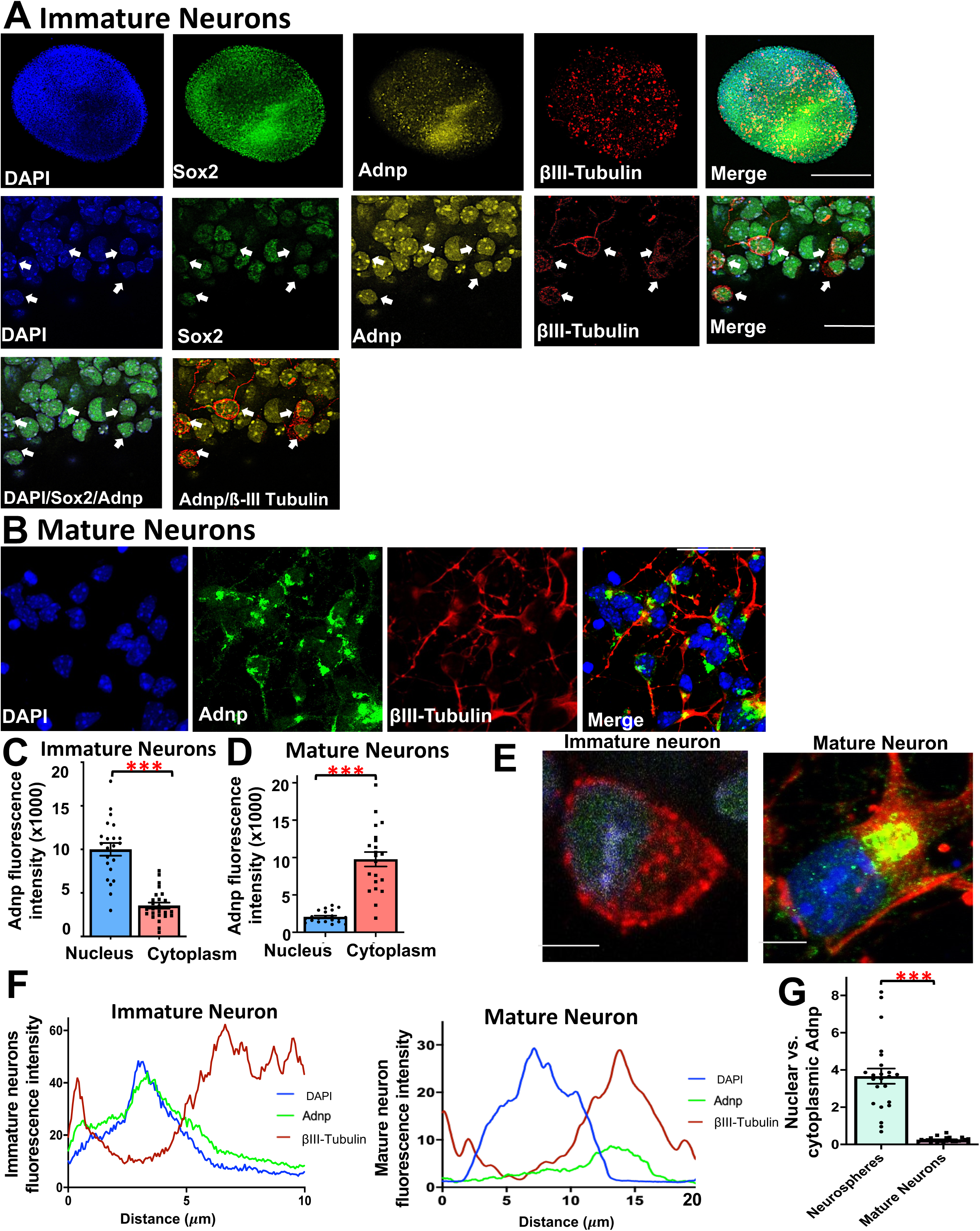
Adnp localization shifts from the nucleus to the cytoplasm as cortical neurons mature. A) Representative photos of Adnp localization in neuronal stem cells grown as neurospheres from low magnification to high magnification. Scale bars = 1 mm (low magnification) and 25 µm (high magnification) B) Representative photos of Adnp localization in primary cortical neurons after neurite elongation from low magnification to high magnification. Scale bars= 50 µm (low magnification) and 25 µm (high magnification) C) Adnp fluorescence intensity in the nucleus and the cytoplasm of neurospheres. Each measurement from the nucleus and the cytoplasm was taken from the same cell. Therefore, a paired samples t-test was used for analysis. This test showed that there is significantly greater Adnp fluorescence in the nucleus (10008 ± 737.4) than in the cytoplasm (3551± 348.9), (95% CI −8045 to −4909), t(21) = 8.589, p <0.001, n=22. D) Adnp fluorescence intensity in the nucleus vs. the cytoplasm of mature neurons. Each measurement was taken from the same cell, therefore a paired-samples t-test was used for analysis. There is significantly greater Adnp fluorescence intensity in the cytoplasm (97730 ± 9555) compared to the nucleus (20705 ± 1694), (95% CI 58088 to 95962), t(20) = 8.484, p<0.0001, n=21. E) High magnification representative photos of the soma of an immature vs. mature neuron. Scale bars= 10 µm, F) Fluorescence intensity plots of the representative cells. G) Ratios of nuclear to cytoplasmic Adnp fluorescence intensity in neurospheres vs. mature neurons. This ratio was calculated by dividing the nuclear fluorescence by the cytoplasmic fluorescence for each cell. An independent samples t-test showed that neurospheres had a significantly higher ratio (3.665 ± 0.408, n=23) than mature neurons (0.247 ± 0.0267, n=24), (95% CI −4.224 to −2.611), t(45) = 8.533, p<0.001. (independent experiments per condition from 3 different litters, stem cells analyzed from 5 distinct neurospheres).

### 14-3-3 inhibition traps Adnp in the nucleus and Adnp binds 14-3-3ε

Next, we investigated the mechanism for Adnp’s nuclear-cytoplasmic shuttling. 14-3-3 proteins are important for development, neurite formation, and are well-known nuclear-cytoplasmic shuttles (8, 41, 54, 55). We performed *in silico* sequence analysis which revealed a likely interaction between Adnp and 14-3-3. To test whether 14-3-3 proteins are involved in Adnp nuclear-cytoplasmic shuttling, we harvested primary cortical neurons at E15.5 and nucleofected them to express either a global 14-3-3 isoform inhibitor, difopein, or the control backbone plasmid, and fixed neurons 48 hours after re-plating and immunofluorescently stained for Adnp (Fig. 11A-B). Performing fluorescence profile analysis of all fluorophores and Adnp fluorescence intensity quantification based on cellular compartment, we found that EYFP-negative control expressing neurons had Adnp fluorescence in the nucleus, but significantly more Adnp fluorescence in the cytoplasm (Fig 11A). Due to difopein expression, Adnp was significantly more localized to the nucleus with a pattern more closely resembling neuronal stem cells than mature neurons (Fig 11B). Because it is well known that 14-3-3ε is important for cortical development and neurite formation (8, 41, 54, 55), we performed pull-down to test 14-3-3ε and Adnp binding. Using COS1 cells, we expressed HAHA-14-3-3ε and 6xHis-Adnp or HAHA and 6xHis-Adnp. We found that Adnp was pulled down by HAHA-14-3-3ε, suggesting that Adnp and 14-3-3ε bind (Fig. 11C). Taken together, these experiments provide mechanistic evidence of Adnp nuclear-cytoplasmic shuttling by 14-3-3ε. We conclude that as neurite formation begins, Adnp is shuttled from the nucleus to the cytoplasm by 14-3-3ε where it promotes neurite formation.

**Figure 11.**
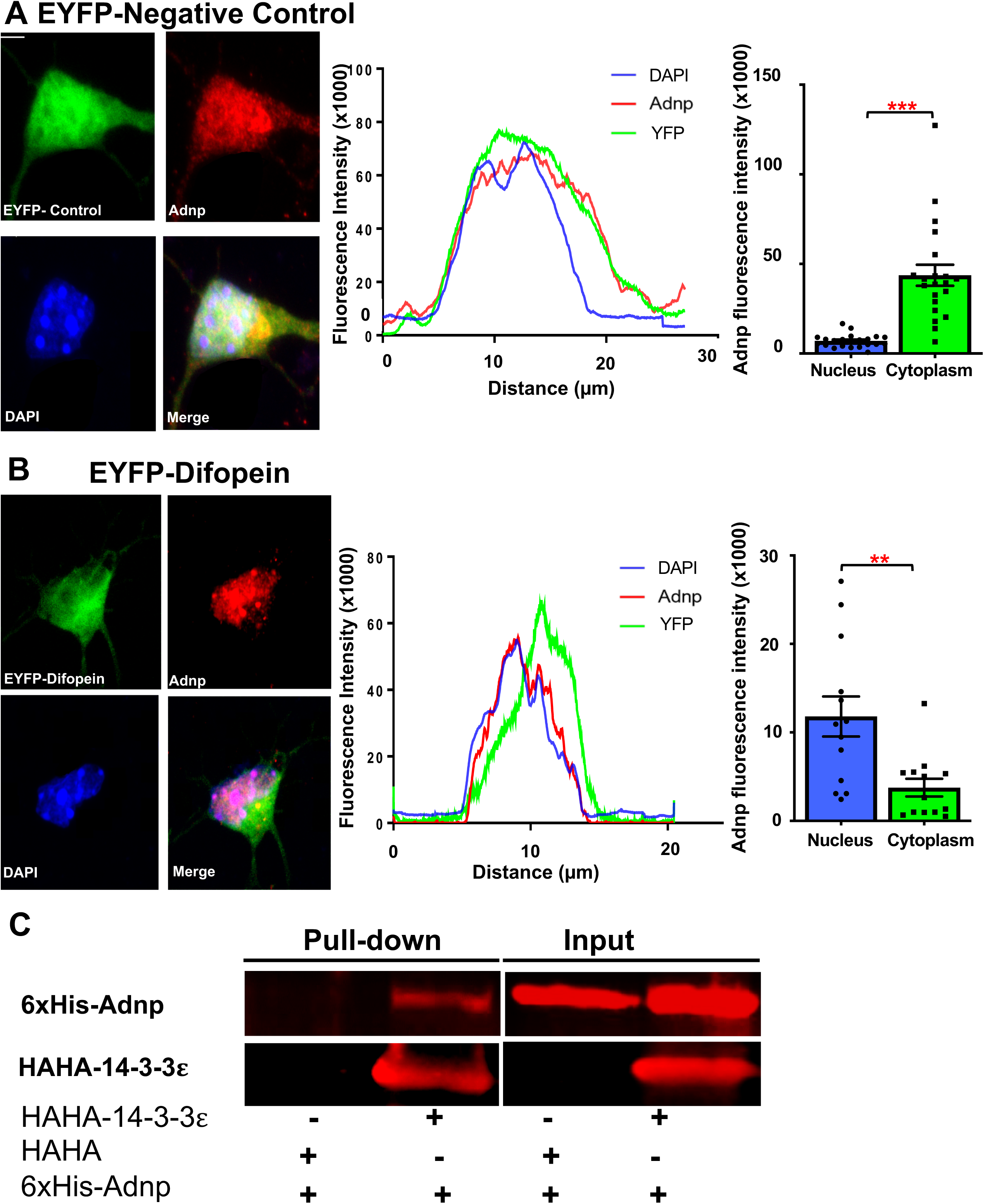
Global 14-3-3 protein inhibition traps Adnp in the nucleus, and Adnp binds 14-3-3*ε*. A) Representative photo of Adnp staining in neurons expressing the backbone plasmid expressing EYFP as a negative control. Scale bar=5 µm. Accompanying the photo is its fluorescence intensity profile for all fluorophores. A paired samples t-test revealed significantly more Adnp fluorescence in the cytoplasm (53563± 8742) compared to the nucleus (7140 ± 763.7), (95% CI 28873 to 63974), t(22) = 5.486, p<0.0001, n=23. B) Representative photo of Adnp staining in neurons expressing EYFP-Difopein which is a global 14-3-3 inhibitor. Scale bar= 5 µm. Accompanying the photo is its fluorescence intensity profile for all fluorophores. A paired samples t-test revealed significantly more Adnp fluorescence in the nucleus (11809 ± 2263) compared to the cytoplasm (3756± 1004), (95% CI −12828 to −3279), t(12) = 3.675, p=0.0032, n=13. (3 independent experiments per condition from 3 different litters). C) Western blot from pull-down analysis using HAHA-conjugated beads, His-tagged Adnp, HAHA-tagged 14-3-3*ε*, and HAHA-tag alone as a negative control. His-Adnp is detected by western blot only when combined with HAHA-14-3-3*ε* and not HAHA, indicating that Adnp is pulled down by 14-3-3*ε*. This suggests that Adnp and 14-3-3*ε* bind. (3 independent experiments).

## Discussion

Changes in expression levels of ADNP is a characteristic of many disorders such as ADNP syndrome, ASD, ID, epilepsy, fetal alcohol syndrome, schizophrenia, and Alzheimer’s disease (23, 30, 36, 56–58). Mutations in the ADNP syndrome patient population are *de novo* heterozygous nonsense and frameshift truncating mutations leading to different outcomes for the expression of the protein depending on the patient population, cell-type, and mutation-type examined. ADNP heterozygous deficiency has been well modeled in the Adnp^+/-^ mouse, which has both behavioral and anatomical defects akin to ADNP syndrome (34). Similarly, somatic mutations in *ADNP* are linked to neurodegeneration in Alzheimer’s disease (25). We performed a cell-type and developmental timing specific analysis of how loss of Adnp affects a specific subpopulation of neurons, layer 2/3 pyramidal neurons, in the developing mouse somatosensory cortex. This cell type was selected because layer 2/3 neurons are essentially integrated into many cortical circuits which are frequently disrupted in patients with ASD, and the location was selected because patients with ADNP syndrome have symptoms that indicate somatosensory processing deficits (23, 59–61). Interestingly these neurons are also possibly vulnerable to aging processes, providing a possible link to Adnp’s roles in neurodegeneration (62). Previous studies that have shown NAP and Adnp are positive regulators of neuritogenesis based on MAP2 fluorescence intensity (31, 37), however, the systems used and study purposes are different than ours. The original Adnp study was looking at neurodifferentiation from multipotent cells, here our initial *in vitro* results, which did not test NAP but only Adnp, indicated an intricate role for Adnp as outlined below. Our results were further confirmed *in vivo*, which was the first time such an in-depth morphological analysis on neurons deficient in Adnp has been performed.

To investigate how loss of Adnp affects neurite formation *in vivo* in real time, we performed *ex vivo* time-lapse live imaging at the onset of neuritogenesis at P0. By P0, the majority of layer 2/3 pyramidal neurons have completed migration and arrived in their final destination at the cortical plate, where they begin neurite initiation followed by neurite elongation. Our results show striking differences between Adnp knockdown and control neurons at this time point. Control neurons are incredibly dynamic, rapidly extending and retracting various processes that are eventually stabilized to elongate. Knockdown neurons slowly but steadily extend one large primary neurite. At the beginning of live imaging, Adnp deficient neurons already had long primary neurites, suggesting that neurons begin neurite formation earlier than control. We looked at earlier developmental time points E17.5 and E18.5 and found no deficits in neuronal migration or morphology. This suggests that both control and Adnp deficient layer 2/3 pyramidal neurons arrive in the cortical plate simultaneously. It is likely that control and Adnp shRNA neurons begin neurite formation simultaneously, but Adnp deficient neurons extend neurites that slowly and stably continue to elongate due to a lack of retraction, whereas control neurons rapidly form and retract many different neurites. This is in agreement with Adnp’s direct involvement in microtubule dynamics (25, 63). However, a direct investigation of how loss of Adnp effects the microtubule network during this developmental process will be crucial to understanding the cellular etiology of ADNP syndrome.

In terms of neurite initiation, Adnp deficient neurons do not extend many, if any, secondary neurites during this imaging. However, when a secondary neurite did emerge it was almost always stabilized, 83% of the time never retracting and re-emerging as is common in control neurons which had a 50% stabilization rate. This result can be attributed to microtubule stabilization and invasion (8, 11–13, 53), in which Adnp has a known role in both the axon growth cone and in developing dendritic spines (25, 63), and we propose that Adnp also fulfills this role during neurite initiation. Taken together the steady growth rate, lack of neurite retraction, and increased neurite stabilization in Adnp deficient neurons are mechanisms that could contribute to the increased basal dendrite number and increased axon length seen in mature Adnp deficient cortical neurons. A final deficit present in these neurons is the emergence of swellings along the primary neurite. These swellings may indicate disrupted intracellular transport of large organelles such as Golgi or mitochondria which typically accumulate in newly emerged dendrites (64). Adnp also has an important known role in microtubule driven intracellular transport of cargo, specifically in the axon (36), but it is possible that Adnp may play a similar role in developing dendrites.

Lastly, we confirmed results present in the Adnp haploinsufficient mouse of decreased dendritic spine density on the basal and apical dendrites of layer 2/3 pyramidal neurons deficient in Adnp (34). Furthermore, we expanded on previous results by performing an unbiased, mathematical quantification of dendritic spine morphology. We found on Adnp deficient dendrites, there are significantly more thin-type spines and significantly fewer mushroom spines compared to control neurons. At P15, many spines adopt a mushroom morphology which can be considered mature and synaptically active (16, 17). These results suggest that not only do Adnp deficient neurons have less spines, but the spines present may not be synaptically active. These results combined with our morphological data presented many possibilities for the activities of these cells.

The increased axon length seen in our Adnp knockdown neurons was a striking and exciting phenotype. Not only are callosally projecting axons from layer 2/3 pyramidal neurons longer, but they innervate much more of the cortex compared to our control axons. We concluded that this phenotype coupled with an increase in basal dendrites could lead to cortico-cortical hyperconnectivity and excitability which is frequently seen in patients with ASD, especially those that also experience seizures as with ADNP syndrome (65–67). This phenotype could be a result of both increased axon length and aberrant axon guidance, which ultimately relies on cytoskeletal dynamics in the axon growth cone. NAP, a short peptide derived from ADNP, is known to regulate microtubule invasion into the axon growth cone (53), so it is logical to suggest that full-length Adnp also regulates this process. However, as with our neurite formation results, NAP was a positive regulator of microtubule invasion (53), whereas our results suggest that full-length Adnp has a more complicated role. This possibility should be thoroughly investigated and provides further evidence for Adnp’s involvement in neuronal morphogenesis being based in a cytoplasmic, microtubule-based mechanism. Another morphological aspect of Adnp deficient neurons is the deviation in the angle of the apical dendrite. Neuronal network formation heavily relies on morphological details such as this, and even small deviations in this angle can have large consequences on functional connectivity (48).

To test the possible effects on neuronal activity suggested by our morphological data, we performed calcium imaging as a measure of neuronal function. Adnp regulates proteins involved in calcium signaling in ways that may affect neuronal plasticity, long-term potentiation, and neurotransmitter release at glutamatergic synapses (63, 68). Calcium signaling is a reliable measure of neuronal function because action potentials trigger large, rapid changes in cytosolic calcium (69). We were the first to assess if Adnp directly regulates calcium signaling. We found increased spontaneous excitability of layer 2/3 pyramidal neurons deficient in Adnp, indicating that the decreased spine density and more immature morphology did not negatively impact the spontaneous activity of these cells. On the contrary, the increased excitability suggests that the increased basal dendrite number may be sufficient to influence neuronal excitability despite the noted spine defects. We attribute the increased calcium signaling to the morphology, as opposed to changes in calcium channel expression, because previous work has actually shown decreases in voltage-gated calcium channel protein expression, Cacnb1, in the hippocampus due to Adnp haploinsufficiency (63). Decreased calcium channel expression would usually result in decreased calcium signaling, but we propose that the increase in dendritic number is again sufficient to overcome this channel deficit as with the dendritic spines. It would be important to investigate this possibility in the cortex, as previous work was performed in the hippocampus, to clarify the mechanism behind changes to calcium signaling in Adnp deficient neurons.

To further dissect how Adnp deficient neurons have increased excitability despite the noted spine defects, we wanted to probe whether cortical connectivity is also disrupted due to Adnp knockdown. Based on our axon tracing analysis which showed increased axon innervation to the opposing cortical hemisphere, we chose to test interhemispheric cortico-cortical connectivity between layer 2/3 pyramidal neurons. Furthermore, many ASD patients have disrupted cortico-cortical connectivity, but there is little consensus in the human population regarding the directionality of these changes (65–67, 70). We utilized the novel technique GRAPHIC to quantify synapses from axons projecting from layer 2/3 pyramidal neurons in the opposing cortical hemisphere. We found a significant increase in GFP puncta on the basal dendrites of Adnp deficient cells. We chose to analyze basal dendrites because these morphological changes of increased number are more likely to contribute to increased connectivity as opposed to the changes in the apical dendrite. The apical dendrite changes observed regarding the angle of extension are more likely to change to type of connectivity as opposed to the amount of connectivity. In addition to increased GFP puncta, we also found that the majority of the puncta were on the dendritic shaft of Adnp deficient neurons as opposed to on the spines as in control samples. This suggests that the immaturity and decreased spine density of Adnp deficient neurons do not negatively affect excitatory connectivity, because the connections being formed are mainly not through dendritic spines. This is similar to an observation in the *Adnp ^+/-^* mouse which showed increased shaft PSD-95 density compared to control animals (34). This phenomenon could contribute to increased excitability as seen in our calcium imaging in multiple ways. Firstly, simply the increase in excitatory contacts from the opposing cortical hemisphere. Secondly, these contacts are on the dendritic shaft, which is the site of approximately 70% of inhibitory contacts (71–73). It is possible that the location of these contacts also contributes to a subsequent decrease in inhibitory synaptic input. It is often noted in cases of ASD that there is a shift in the excitatory/inhibitory balance of neurons and circuits, and our results suggest that this is also the case for ADNP syndrome. The current study only assesses interhemispheric excitatory connections, but our morphological assessment suggests that an increase in intrahemispheric connections is also highly likely. Future studies should aim to clarify intrahemispheric connectivity as well as inhibitory connectivity to gain a comprehensive understanding of how cortical connectivity is altered due to loss of Adnp. The implications these results have on functional cortical connectivity, potentially of multiple cortical circuits, cannot be understated.

To probe the appropriate cellular mechanism underlying our morphological phenotypes we first performed Adnp localization analysis as neurons underwent neuritogenesis, because Adnp is a multifunctional protein with roles in both the nucleus and the cytoplasm (22, 23, 34, 35, 53, 74). We found that Adnp is shuttled from the nucleus to the cytoplasm in neuronal stem cells versus neurons in late-stages of neurite elongation, 48 hours following re-plating. It has been previously reported that Adnp is crucial for stem cell and embryo development (51, 52, 75, 76), and upon retinoic acid induced differentiation of embryonic carcinoma cells (P19) Adnp is shuttled from the nucleus to the cytoplasm (37). Our study confirms these results and is the first to show this phenomenon in a neuronal model. These results suggest that Adnp changes its function from mostly nuclear to remodel chromatin and promotes expression of neuronal differentiation inducing genes to mostly cytoplasmic to promote neurite formation. Furthermore, our *in vivo* staining patterns reveal Adnp is located exclusively in the cytoplasm of neurons in the cortex at P15, which is when our morphological analyses took place. 14-3-3 proteins are well known nuclear-cytoplasmic shuttles that are crucial for neuronal development and neurite formation (8, 41, 55, 77–79). By expressing a global 14-3-3 isoform inhibitor in primary cortical neurons we trapped the majority of Adnp in the nucleus, resulting in cells with Adnp localization that was similar to that of neuronal stem cells. Some Adnp was still present in the cytoplasm of these neurons, possibly due to our use of a global inhibitor as opposed to a specific isoform knockdown. We performed *in silico* sequencing analysis that revealed a likely interaction between 14-3-3ε and Adnp, and we tested this by performing a pull-down experiment where we found that Adnp and 14-3-3ε do bind. This mechanism requires further study to identify the binding site and upstream signals, such as kinases, that promote this timing specific interaction and subsequent shuttling. This shuttling mechanism could also be important for understanding cases of ADNP syndrome where mutations result in a nuclear/cytoplasmic localization shift in ADNP, as opposed to a decrease in protein expression (80).

Taken together, this information allows us to propose a model for how Adnp expression and subcellular localization changes to promote neurite formation during cortical development: Adnp is highly localized in the nucleus in immature neurons to regulate the expression of lineage-specific genes and to promote differentiation (51, 52, 74), then as differentiation begins Adnp travels into the cytoplasm aided by 14-3-3ε and along developing neurites where it leads to appropriate neurite formation (Fig. 12A). When expression of Adnp is decreased in layer 2/3 pyramidal neurons, as in ADNP syndrome and other developmental and psychiatric disorders, neurite formation is increased resulting in a longer axon and more basal dendrites and subsequent increased spontaneous excitability and interhemispheric cortico-cortical connectivity (Fig 12B). Further mechanistic studies for how Adnp promotes proper neuronal morphology once it is present in the cytoplasm will be crucial to gain a proper understanding of the cellular etiology of how loss of ADNP effects this developmental process. Our results reveal a snowball-effect of complex morphological and functional deficits in layer 2/3 pyramidal neurons in the developing mouse somatosensory cortex due to loss of Adnp, and help shed light on the phenotypic complexity seen in both the Adnp haploinsufficient mouse as well as the ADNP syndrome patient population (23, 34). We have pinpointed that P0 is when deficits begin for Adnp deficient neurons and these severe deficits are sustained and worsened throughout development; seeming to affect many subsequent maturation stages such as axon guidance, calcium signaling, and cortical connectivity. Future studies to uncover the mechanism behind the primary phenotype, which we pinpointed as neuritogenesis, will be crucial for therapeutic innovation, including NAP potential homeostatic activity.

**Figure 12.**
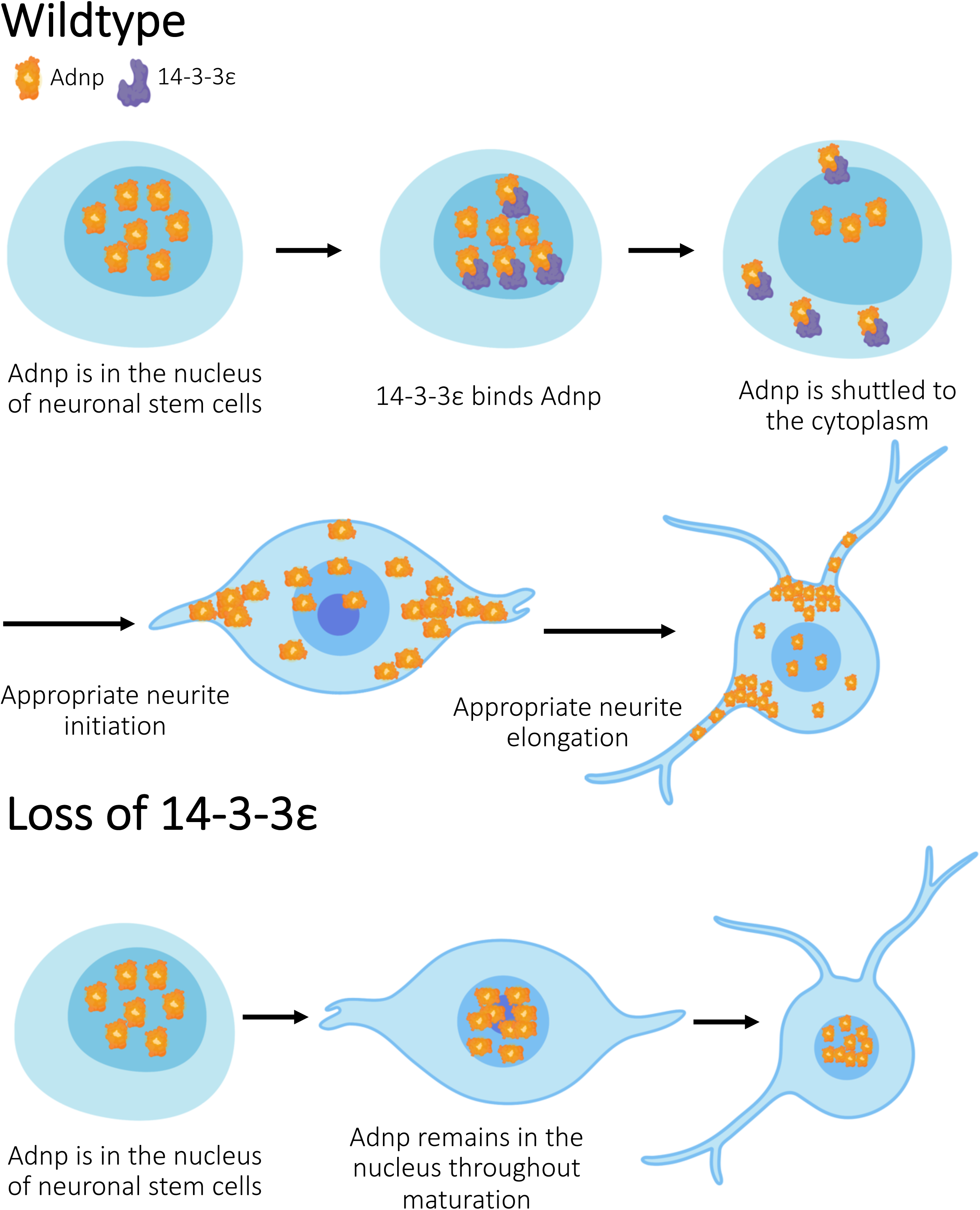
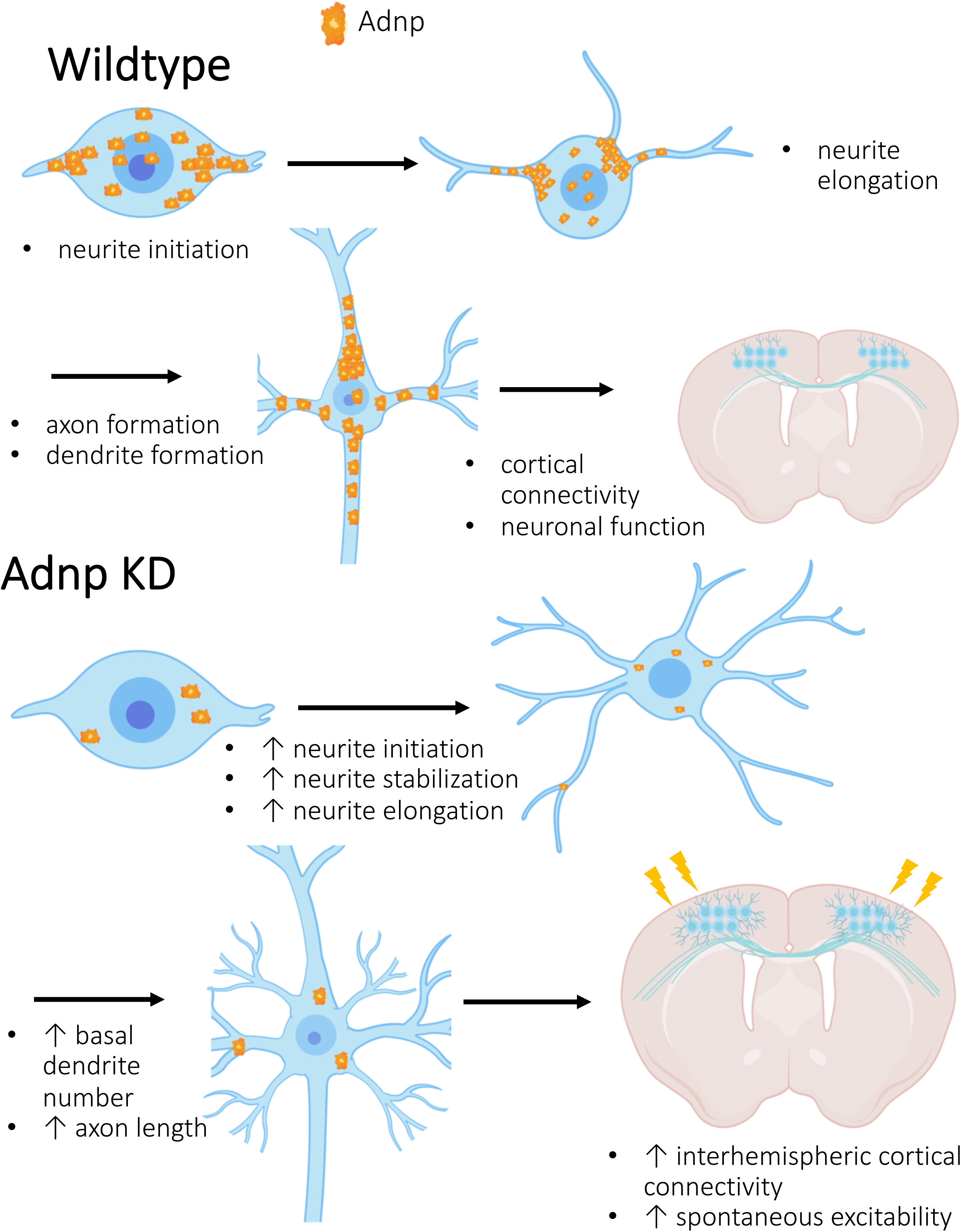
Model for Adnp’s regulation of neurite formation. A) Adnp is present in the nucleus of neuronal stem cells. Upon differentiation, Adnp binds to and subsequently shuttled out of the nucleus by 14-3-3*ε* where it promotes appropriate neurite formation. Proper neurite formation allows for proper axon and dendrite development, which ultimately results in proper neuronal connectivity and function. B) Partial loss of Adnp results in increased neurite initiation of neurites likely to become dendrites, and increased neurite elongation of the neurite likely to become the axon. This results in increased basal dendrite formation and axon length, which ultimately results in increased spontaneous cortical excitability. Figures created using biorender.com.

## Acknowledgements

We would like to thank Dr. Peter Baas, Dr. Wen-Jun Gao, Dr. Elias Spiliotis, Dr. Pat Levitt and Dr. Illana Gozes for their helpful comments, feedback, and fruitful discussions on this manuscript. We also thank Dr. Tomomi Shimogori in Riken Center for Brain Science in Japan who provided the GRAPHIC vectors.

This work has been supported by a research grant from the NINDS (NS096098).

---

**Supplemental Figure 1. Adnp shRNA can effectively knockdown Adnp.**

A-B) Representative photos of endogenous Adnp staining in mature primary cortical neurons expressing Scramble shRNA or Adnp shRNA at low magnification (A) and high magnification (B). C) Comparison of Adnp fluorescence intensity in Scramble shRNA and Adnp shRNA expressing neurons. An independent samples t-test shows that Adnp shRNA expressing neurons have a significantly lower Adnp fluorescence intensity (72290 ± 16469) compared to Scramble shRNA (599955 ± 61742), (95% CI −656146 to −399184), t(48) = 8.258, p<0.001. (n=25, 3 independent experiments per condition from 3 different litters). D) Representative western blots validating our Adnp shRNA, Adnp-His, and scramble shRNA plasmids used in all experiments. Plasmids were co-transfected into HEK293T cells. Quantification of westerns blots which were performed in triplicate. Adnp expression was normalized to total protein expression in each sample using REVERT total protein stain. An independent samples t-test showed that Adnp shRNA significantly reduced Adnp-His protein expression (3.480 x 10^−3^ ± 2.497 x 10^−3^) compared to Scramble shRNA (0.128 ± 0.048), (95% CI −0.2289 to −0.01994), t(5) = 3.061, p = 0.0281, with ∼ 97% efficiency. E) Representative western blots validating our Scramble shRNA, Adnp shRNA, and shRNA Resistant Adnp (Resist Adnp-His) plasmids used in rescue experiments. Plasmids were co-transfected into HEK293T cells. Quantification of western blots which were performed in triplicate. Adnp expression was normalized to total protein expression in each sample using REVERT total protein stain. An independent samples t-test showed that there was no significant difference in Adnp expression between Adnp shRNA and Scramble shRNA when co-expressed with Resist Adnp, p=0.443.

**Supplemental Figure 2. Shorter neurites’ length is not affected by Adnp KD *in vitro*.**

Quantification of length of the shorter neurites, as opposed to the longest neurite. A one-way ANOVA showed that there were no significant differences between groups, F(3, 149) = 2.146, p=0.0968. (Scramble shRNA n=37, Adnp shRNA n=40, Rescue n=38, Rescue control n=38, 3 independent experiments per condition from 3 different litters)

**Supplemental Figure 3. IUE targets neurons that migrate to layer 2/3.**

IUE of mouse embryos at E15.5 specifically transfects layer 2/3 neurons. The brain was harvested at P15. Brn2 staining was used because it specifically marks layers 2/3 and 5 (47). A Scramble-shRNA-Venus plasmid was used for electroporation. There is clear colocalization of Venus and Brn2. Scale bar= 50 µm.

**Supplemental Figure 4. Apical dendrite length is not affected by Adnp KD *in vivo*.**

Quantification of apical dendrite length at P15 after IUE at E15.5 using a one-way ANOVA revealed no significant differences between groups, F(3, 93) = 0.3682, p=0.776, (Scramble shRNA n=25, Adnp shRNA n=24, Rescue n=25, Rescue control n=23, 3 independent experiments per condition from 3 different litters)

**Supplemental Figure 5. Apical dendrite deficits from P3 are sustained through P15.**

A-D) IUE was performed at E15.5 and brains were collected at P15. Representative photos of layer 2/3 pyramidal neurons expressing Scramble shRNA (A), Adnp shRNA (B), rescue plasmids Adnp-shRNA-Venus and Adnp-mScarlet (“Adnp OE”) (C), and rescue control plasmids Scramble-shRNA-Venus and mScarlet backbone vector (“Control OE”) (D). E-H) Polar histograms of Scramble shRNA (E), Adnp shRNA (F), rescue (G), and rescue control (H) expressing neurons which quantifies the angles at which the apical dendrites extended with respect to the cortical plate. I) Quantification of the width of the apical dendrites with respect to the soma. A one-way ANOVA showed that there was a significant difference between groups, F(3,89) = 13.263, p<0.0001, n=24 Bonferroni post hoc comparison showed that Adnp shRNA neurons had significantly wider apical dendrites with respect to the soma (0.239*μ*m ± 0.014*μ*m) compared to Scramble shRNA (0.156*μ*m ± 0.012*μ*m, p<0.0001), rescue (0.143*μ*m ± 0.011*μ*m, p<0.0001), and rescue control neurons (0.161*μ*m ± 0.011*μ*m, p<0.0001). Scramble shRNA neurons did not significantly differ from the rescue neurons (p=1.000) or the rescue control neurons (p=1.000). J) Quantification of the angle of the apical dendrite from the soma. A one-way ANOVA showed that there was a difference in apical dendrite angle deviation between groups F(3,112)= 6.457, p<0.0001, n=29. Bonferroni post hoc comparison revealed that Adnp shRNA expressing neurons had a significantly greater deviation from 90° (18.061° ± 1.452°) compared to Scramble shRNA (10.924° ± 1.147°, p=0.002), rescue (11.441° ± 1.405°, p=0.004), and rescue control (11.303° ± 1.369°, p=0.003). Scramble shRNA neurons did not significantly differ from the rescue neurons (p=1.000) or the rescue control neurons (p=1.000). Scale bars= 100*μ*m. (3 independent experiments per condition from 3 different litters).

**Supplemental Figure 6. Adnp KD neurons have no cortical distribution defects following neurogenesis or neuronal migration.**

MZ= marginal zone, CP= cortical plate, IZ= intermediate zone, SVZ/VZ= subventricular zone/ventricular zone. A) Representative photos of E17.5 cortices after IUE at E15.5, just following neurogenesis. According to a two-way repeated measures ANOVA, Adnp shRNA neurons had no significant differences in percentage of cells in each cortical zone compared to Scramble shRNA neurons, F(4, 12)= 6.341 x 10^−7^, p=1.000. B) Representative photos of E18.5 cortices after IUE at E15.5. According to a two-way repeated measures ANOVA, Adnp shRNA neurons had no significant differences in percentage of cells in each cortical zone compared to Scramble shRNA neurons, F(4, 12)= 2.193 x 10^−18^, p=1.000. C) Representative photos of P3 cortices after IUE at E15.5. According to a two-way repeated measures ANOVA, Adnp shRNA neurons had no significant differences in percentage of cells in each cortical zone compared to Scramble shRNA neurons F(4, 12)= 8.371 x 10^−17^, p=1.000. Scale bars = 100*μ*m. (3 independent experiments from 3 different litters).

