## Supplemental Figures for "14-3-3 shuttles Activity-dependent neuroprotective protein to the cytoplasm to promote appropriate neuronal morphogenesis, cortical connectivity and calcium signaling"

Bennison et al.

### Supplemental Figure 1.

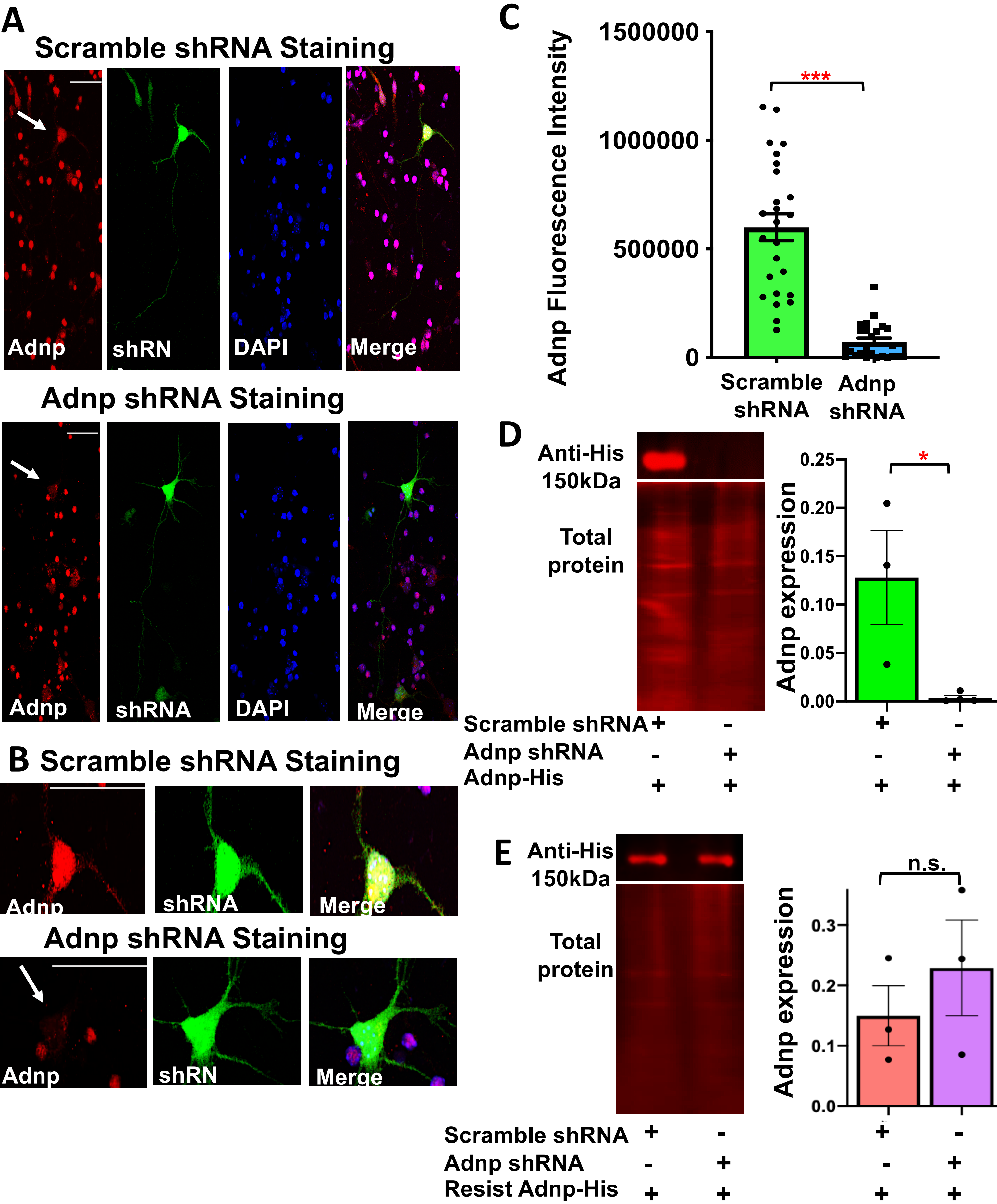

### Supplemental Figure 2.

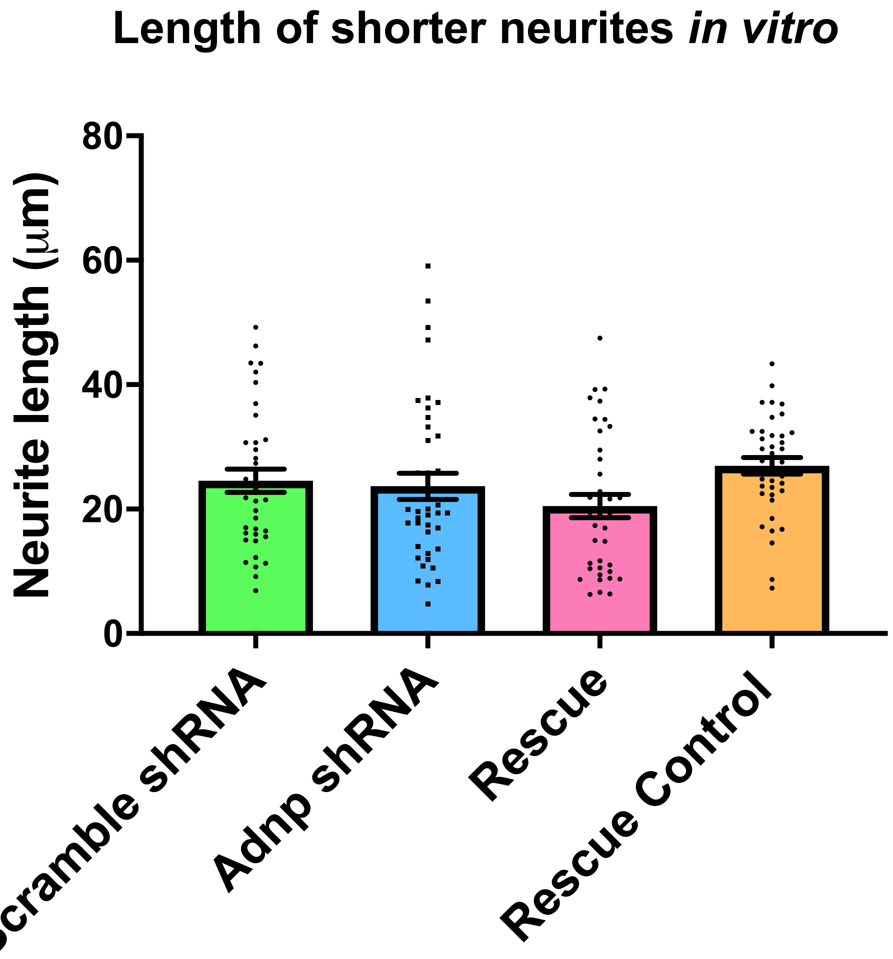

### Supplemental Figure 3.

Bennison et al.

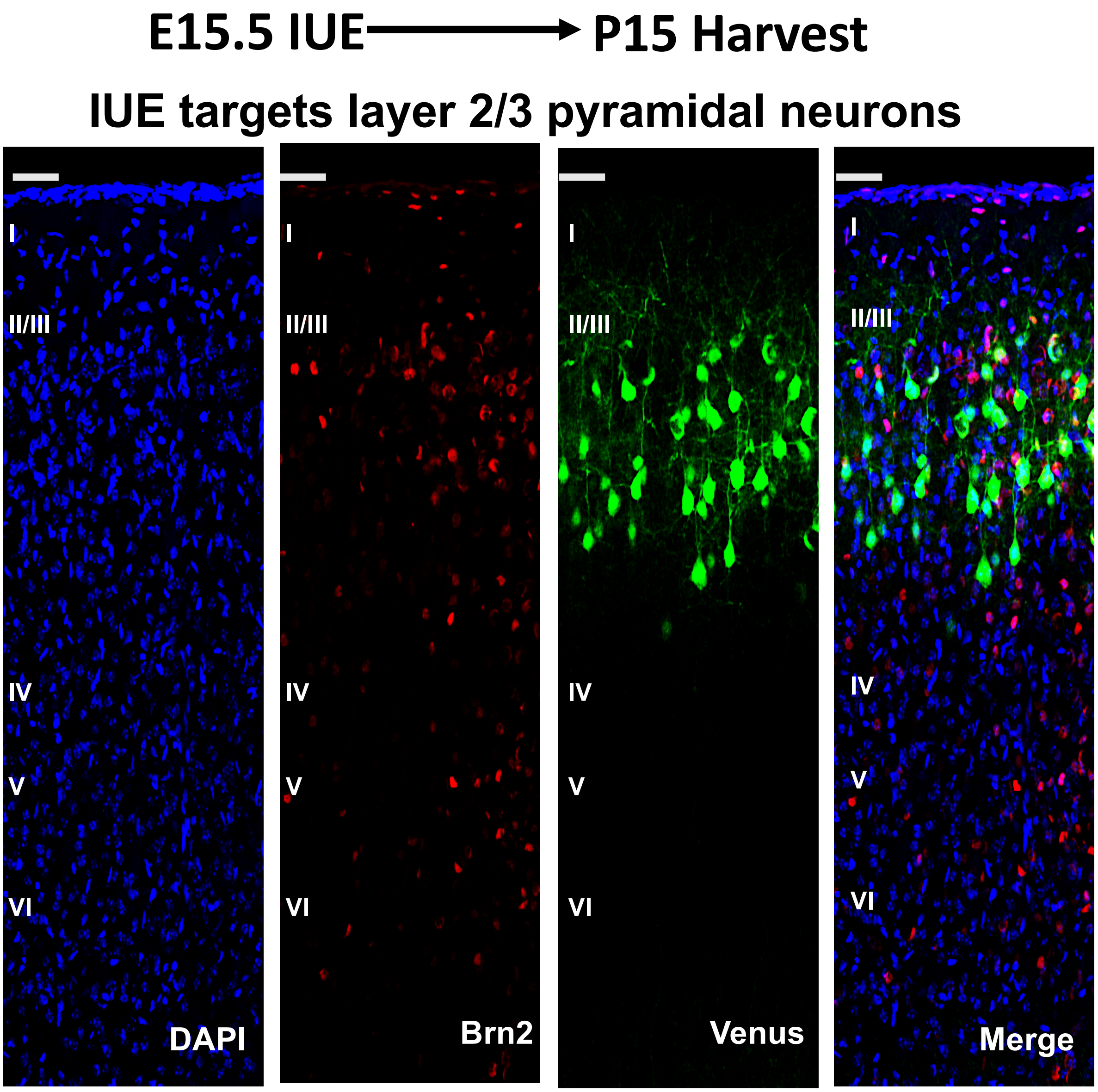

### Supplemental Figure 4.



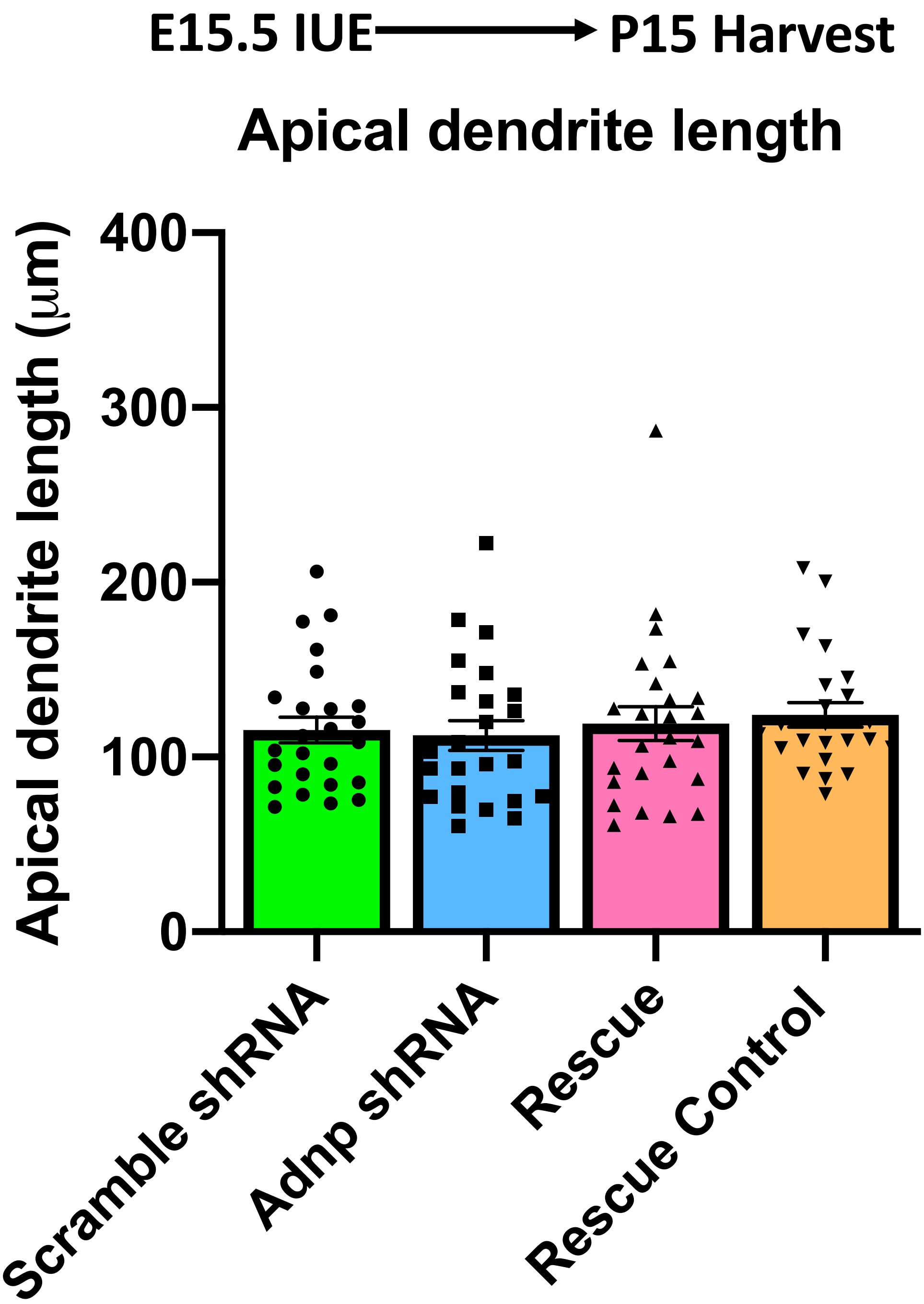

Supplemental Figure 5. Bennison et al.

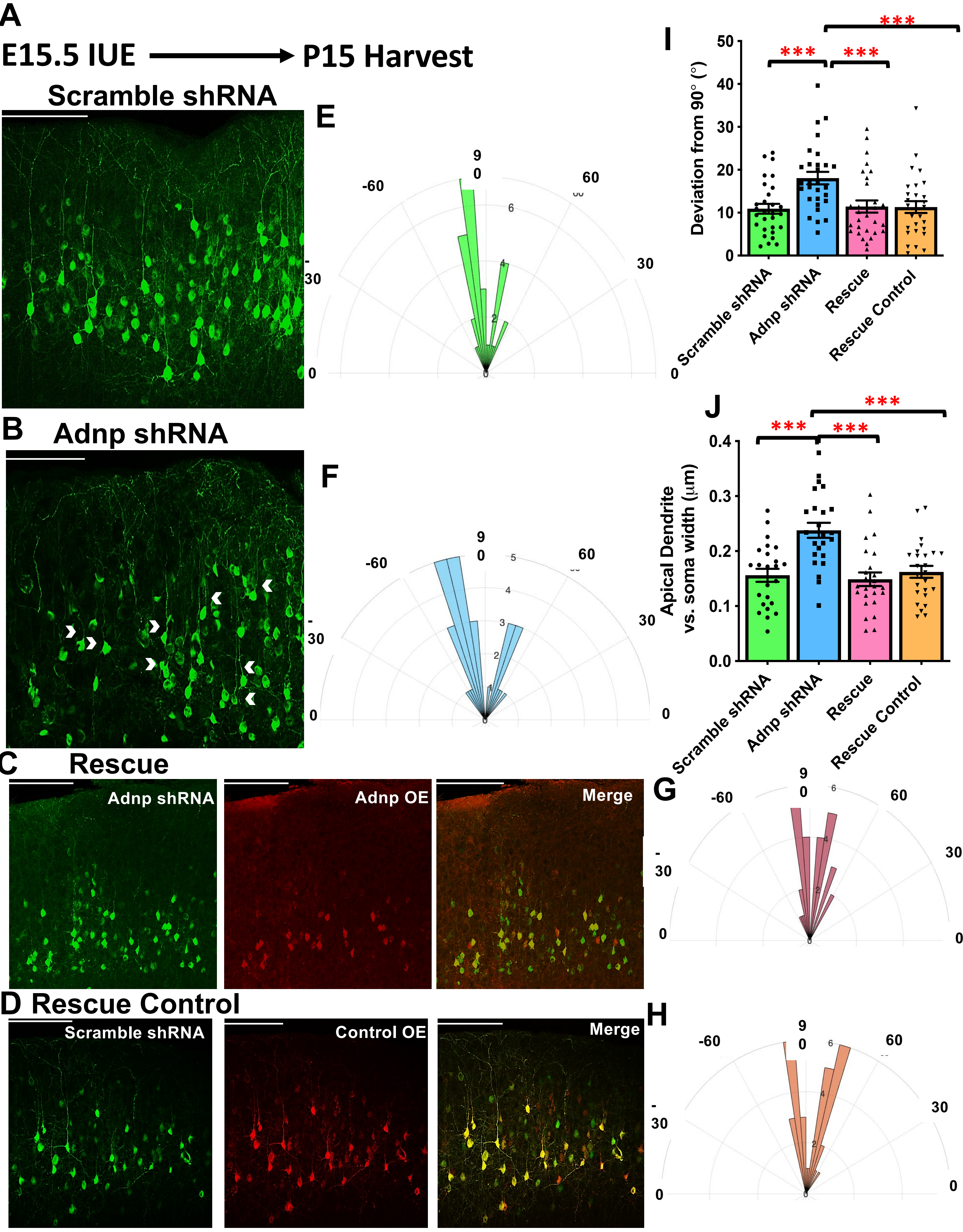

Supplemental Figure 6.

Bennison et al.

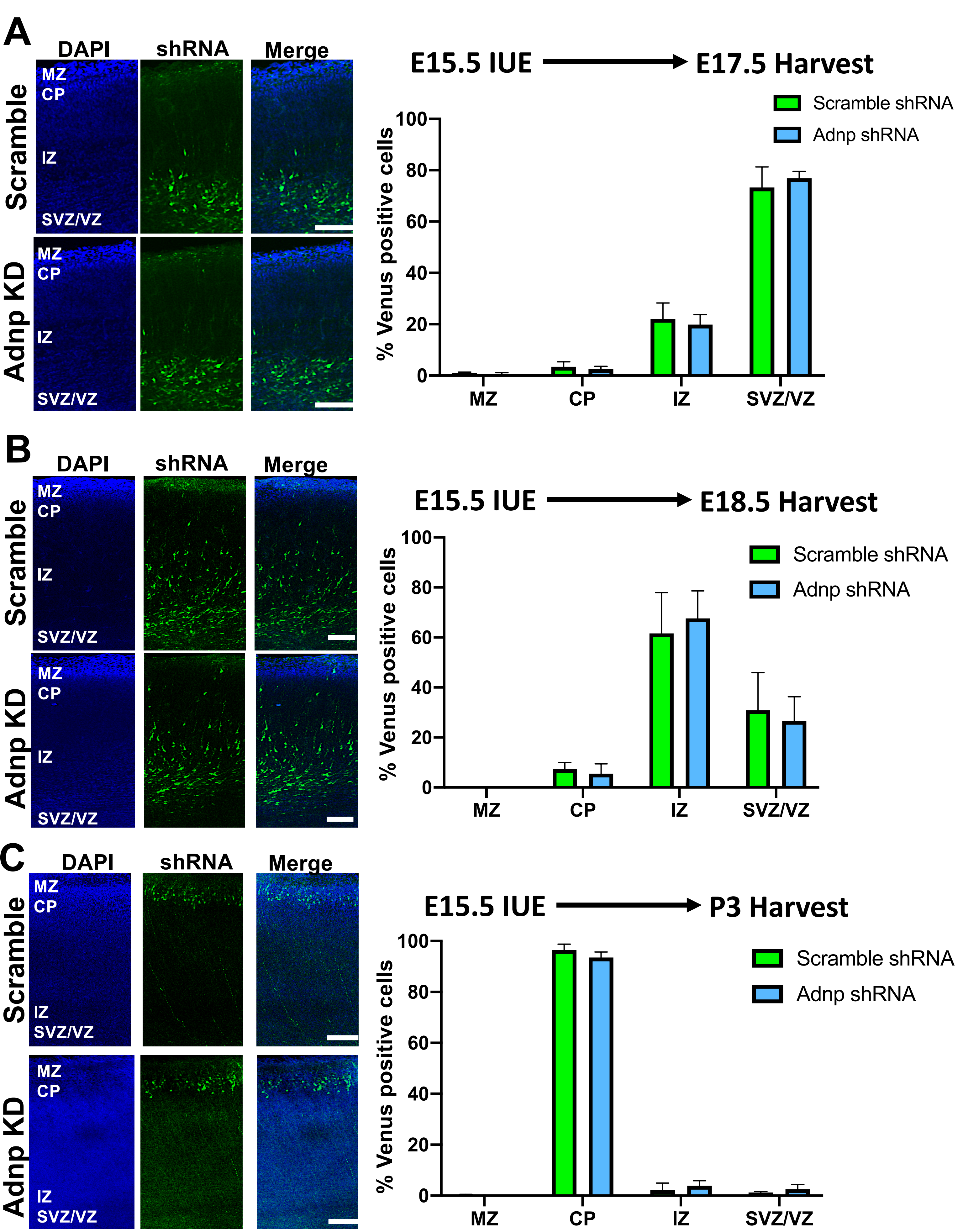
